# Loss of UGP2 in brain leads to a severe epileptic encephalopathy, emphasizing that bi-allelic isoform specific start-loss mutations of essential genes can cause genetic diseases

**DOI:** 10.1101/799841

**Authors:** Elena Perenthaler, Anita Nikoncuk, Soheil Yousefi, Woutje M. Berdowski, Ivan Capo, Herma C. van der Linde, Paul van den Berg, Edwin H. Jacobs, Darija Putar, Mehrnaz Ghazvini, Eleonora Aronica, Wilfred F.J. van IJcken, Walter G. de Valk, Evita Medici-van den Herik, Marjon van Slegtenhorst, Lauren Brick, Mariya Kozenko, Jennefer N. Kohler, Jonathan A. Bernstein, Kristin G. Monaghan, Amber Begtrup, Rebecca Torene, Amna Al Futaisi, Fathiya Al Murshedi, Renjith Mani, Faisal Al Azri, Erik-Jan Kamsteeg, Majid Mojarrad, Atieh Eslahi, Zaynab Khazaei, Fateme Massinaei Darmiyan, Mohammad Doosti, Ehsan Ghayoor Karimiani, Jana Vandrovcova, Faisal Zafar, Nuzhat Rana, Krishna K. Kandaswamy, Jozef Hertecant, Peter Bauer, Stephanie Efthymiou, Henry Houlden, Aida M. Bertoli-Avella, Reza Maroofian, Kyle Retterer, Alice S. Brooks, Tjakko J. van Ham, Tahsin Stefan Barakat

**Affiliations:** Department of Clinical Genetics, Erasmus MC University Medical Center, Rotterdam, The Netherlands; Department for Histology and Embryology, Faculty of Medicine Novi Sad, University of Novi Sad, Serbia; iPS cell core facility, Erasmus MC University Medical Center, Rotterdam, The Netherlands; Amsterdam UMC, University of Amsterdam, Department of (Neuro)pathology, Amsterdam, Amsterdam Neuroscience, The Netherlands; Stichting Epilepsie Instellingen Nederland (SEIN), The Netherlands; Center for Biomics, Department of Cell Biology, Erasmus MC University Medical Center, Rotterdam, The Netherlands; Department of Neurology, Erasmus MC University Medical Center, Rotterdam, The Netherlands; Division of Genetics, McMaster Children’s Hospital, Hamilton, Ontario, L8S 4J9, Canada; Division of Cardiovascular Medicine, Stanford University School of Medicine, Stanford, CA, 94035, USA; Department of Pediatrics, Division of Medical Genetics, Stanford University School of Medicine, Stanford, CA 94035, USA; GeneDx, Gaithersburg, MD, 20877, USA; Department of Child health, college of Medicine and Health Sciences, Sultan Qaboos University, Muscat, Oman; Genetic and Developmental Medicine Clinic, Sultan Qaboos University Hospital, Muscat, Oman; Department of Radiology and molecular imaging, Sultan Qaboos University Hospital, Muscat, Oman; Department of Clinical Genetics, Radboud University, Nijmegen, The Netherlands; Department of Medical Genetics, Faculty of Medicine, Mashhad University of Medical Sciences, Mashhad, Iran; Medical Genetics Research Center, Mashhad University of Medical Sciences, Mashhad, Iran; Genetic Center of Khorasan Razavi, Mashhad, Iran; Student Research Committee, Faculty of Medicine, Mashhad University of Medical Sciences, Mashhad, Iran; Genetic Counseling Center, Welfare Organization of Sistan and Baluchestan, Zahedan, Iran; Department of Modern Sciences and Technologies, Faculty of Medicine, Mashhad University of Medical Sciences, Mashhad, Iran; Genetics Research Centre, Molecular and Clinical Sciences Institute, St. George’s, University, London, SW17 ORE, United Kingdom; Department of Neuromuscular Disorders, UCL Queen Square Institute of Neurology, London, WC1N 3BG, United Kingdom; Department of Paediatric Neurology, Children’s hospital and institute of Child health, Multan 60000, Pakistan; CENTOGENE AG, Rostock, Germany; Department of Pediatrics, Tawam Hospital, and College of Medicine and Health Sciences, UAE University, Al-Ain, UAE

**Keywords:** epileptic encephalopathy, UGP2, ATG mutations, start-loss mutation, genetics, whole exome sequencing, microcephaly, recurrent mutation, founder mutation, essential gene

## Abstract

Developmental and/or epileptic encephalopathies (DEEs) are a group of devastating genetic disorders, resulting in early onset, therapy resistant seizures and developmental delay. Here we report on 12 individuals from 10 families presenting with a severe form of intractable epilepsy, severe developmental delay, progressive microcephaly and visual disturbance. Whole exome sequencing identified a recurrent, homozygous variant (chr2:64083454A>G) in the essential *UDP-glucose pyrophosphorylase* (*UGP2*) gene in all probands. This rare variant results in a tolerable Met12Val missense change of the longer UGP2 protein isoform but causes a disruption of the start codon of the shorter isoform. We show that the absence of the shorter isoform leads to a reduction of functional UGP2 enzyme in brain cell types, leading to altered glycogen metabolism, upregulated unfolded protein response and premature neuronal differentiation, as modelled during pluripotent stem cell differentiation *in vitro*. In contrast, the complete lack of all UGP2 isoforms leads to differentiation defects in multiple lineages in human cells. Reduced expression of Ugp2a/Ugp2b *in vivo* in zebrafish mimics visual disturbance and mutant animals show a behavioral phenotype. Our study identifies a recurrent start codon mutation in *UGP2* as a cause of a novel autosomal recessive DEE. Importantly, it also shows that isoform specific start-loss mutations causing expression loss of a tissue relevant isoform of an essential protein can cause a genetic disease, even when an organism-wide protein absence is incompatible with life. We provide additional examples where a similar disease mechanism applies.

## Introduction

Developmental and/or epileptic encephalopathies (DEEs) are a heterogeneous group of genetic disorders, characterized by severe epileptic seizures in combination with developmental delay or regression^1^. Genes involved in multiple pathophysiological pathways have been implicated in DEEs, including synaptic impairment, ion channel alterations, transporter defects and metabolic processes such as disorders of glycosylation^2^. Mostly, dominant acting, *de novo* mutations have been identified in children suffering from DEEs^3^, and only a limited number of genes with a recessive mode of inheritance are known so far, with a higher occurrence rate in consanguineous populations^4^. A recent cohort study on DEEs employing whole exome sequencing (WES) and copy-number analysis, however, found that up to 38% of diagnosed cases might be caused by recessive genes, indicating that the importance of this mode of inheritance in DEEs has been underestimated^5^.

The human genome contains ∼20,000 genes of which more than 5,000 have been implicated in genetic disorders. Wide-scale population genomics studies and CRISPR-Cas9 based loss-of-function (LoF) screens have identified around 3000-7000 genes that are essential for the viability of the human organism or result in profound loss of fitness when mutated. In agreement with that they are depleted for LoF variants in the human population^6^. For some of these essential genes it is believed that LoF variants are incompatible with life and are therefore unlikely to be implicated in genetic disorders presenting in postnatal life^7^. One such example is the *UDP-glucose pyrophosphorylase* (*UGP2*) gene at chromosome 2. UGP2 is an essential octameric enzyme in nucleotide-sugar metabolism^8-10^, as it is the only known enzyme capable of catalyzing the conversion of glucose-1-phosphate to UDP-glucose^11,12^. UDP-glucose is a crucial precursor for the production of glycogen by *glycogen synthase* (GYS)^13,14^, and also serves as a substrate for *UDP-glucose:glycoprotein transferases* (UGGT) and *UDP-glucose-6-dehydrogenase* (UGDH), thereby playing important roles in glycoprotein folding control, glycoconjugation and UDP-glucuronic acid synthesis. The latter is an obligate precursor for the synthesis of glycosaminoglycans and proteoglycans of the extracellular matrix^15,16^, of which aberrations have been associated with DEEs and neurological disorders^17-20^. *UGP2* has previously been identified as a marker protein in various types of malignancies including gliomas where its upregulation is correlated with a poor disease outcome^21-28^, but has so far not been implicated in genetic diseases and it has been speculated that this is given its essential role in metabolism^8^.

Many genes are differentially expressed amongst tissues, regulated by non-coding regulatory elements^29^. In addition, it has become clear that there are more than 40,000 protein isoforms encoded in the human genome, whose expression levels vary amongst tissues. Although there are examples of genetic disorders caused by the loss of tissue specific protein isoforms^30-33^, it is unknown whether a tissue-relevant loss of an essential gene can be involved in human disease. Here, we report on such a scenario, providing evidence that a novel form of a severe DEE is caused by the brain relevant loss of the essential gene *UGP2* due to an isoform specific and germ line transmitted start codon mutation. We present data that this is likely a more frequent disease mechanism in human genetics, illustrating that essential genes for which organism-wide loss is lethal can still be implicated in genetic disease when only absent in certain tissues due to expression misregulation.

## Results

### A recurrent ATG mutation in UGP2 in 12 individuals presenting with a severe DE

We encountered a three-month old girl (**Figure 1A**, family 1, individual 1), that was born as the first child to healthy non-consanguineous Dutch parents, by normal vaginal delivery after an uneventful pregnancy conceived by ICSI. She presented in the first weeks of life with irritability and jitteriness, that developed into infantile spasms and severe epileptic activity on multiple electroencephalograms, giving rise to a clinical diagnosis of West syndrome **(Figure 1B**). Despite the use of multiple anti-epileptic drugs, including ACTH and a ketogenic diet, seizures remained intractable and occurred daily. Severe developmental delay was evident without acquisition of any noticeable developmental milestones, causing the need for gastrointestinal tube feeding. Visual tracking was absent, and foveal hypopigmentation, hypermetropia and mild nystagmus were noticed upon ophthalmological investigation. MRI brain imaging showed no gross structural abnormalities or migration disorders at the age of 4 months, but displayed reduced white matter, that further developed into global atrophy with wide sulci and wide pericerebral liquor spaces at the age of 17 months (**Figure 1C, Supplementary Figure 1B**). At that time, she had become progressively microcephalic, with a head circumference of −2.96 SD at the last investigation at 23 months of age (**Supplementary Figure 1A**). She showed a number of minor dysmorphisms, including a sloping forehead, elongated head with suture ridging, bitemporal narrowing, a relatively small mouth and large ears (**Figure 1A**). Neurological examination showed brisk, symmetric deep tendon reflexes, more pronounced at the upper limbs. Routine investigations, including metabolic screening in urine, plasma and cerebrospinal fluid were normal. A SNP-array showed a normal female chromosomal profile, with a large, ∼30 Mb run of homozygosity (ROH) at chromosome 2, and a few smaller ROH regions, adding up to 50 Mb ROH regions in total, pointing to an unrecognized common ancestor of both parents (coefficient of inbreeding 1/64). Subsequent trio WES did not show any disease-causing variants in known DEE genes, but identified a homozygous variant (chr2:64083454A>G) in *UGP2*, located in the large ROH region (**Figure 1D**), with no other disease implicated variants observed in that region. Both parents were heterozygous carriers of the same variant. Via Genematcher^34^ and our network of collaborators, we identified 11 additional individuals from 9 unrelated families (of which 8 were consanguineous), harboring the exact same homozygous variant and presenting with an almost identical clinical phenotype of intractable seizures, severe developmental delay, visual disturbance, microcephaly and similar minor dysmorphisms (**Figure 1A, C, E, Supplementary Figure 1B, Supplementary Case reports, Supplementary Table 1** for detailed information on 11 cases). Six of these individuals passed away before the age of 3.5 years. In 4 families, at least 4 already deceased siblings had a similar phenotype but could not be investigated. Two families were of Indian descent (both with ancestors from regions currently belonging to Pakistan), living in Canada (family 2) and the USA (family 3), with the remaining families coming from Oman (family 4, originally from Pakistan), Pakistan (family 5), Iran (family 6, 7 and 8) and UAE (family 9). One additional case in a family from Oman was identified presenting with intractable seizures and microcephaly, but no detailed medical information could be obtained at this point.

**Figure 1:**
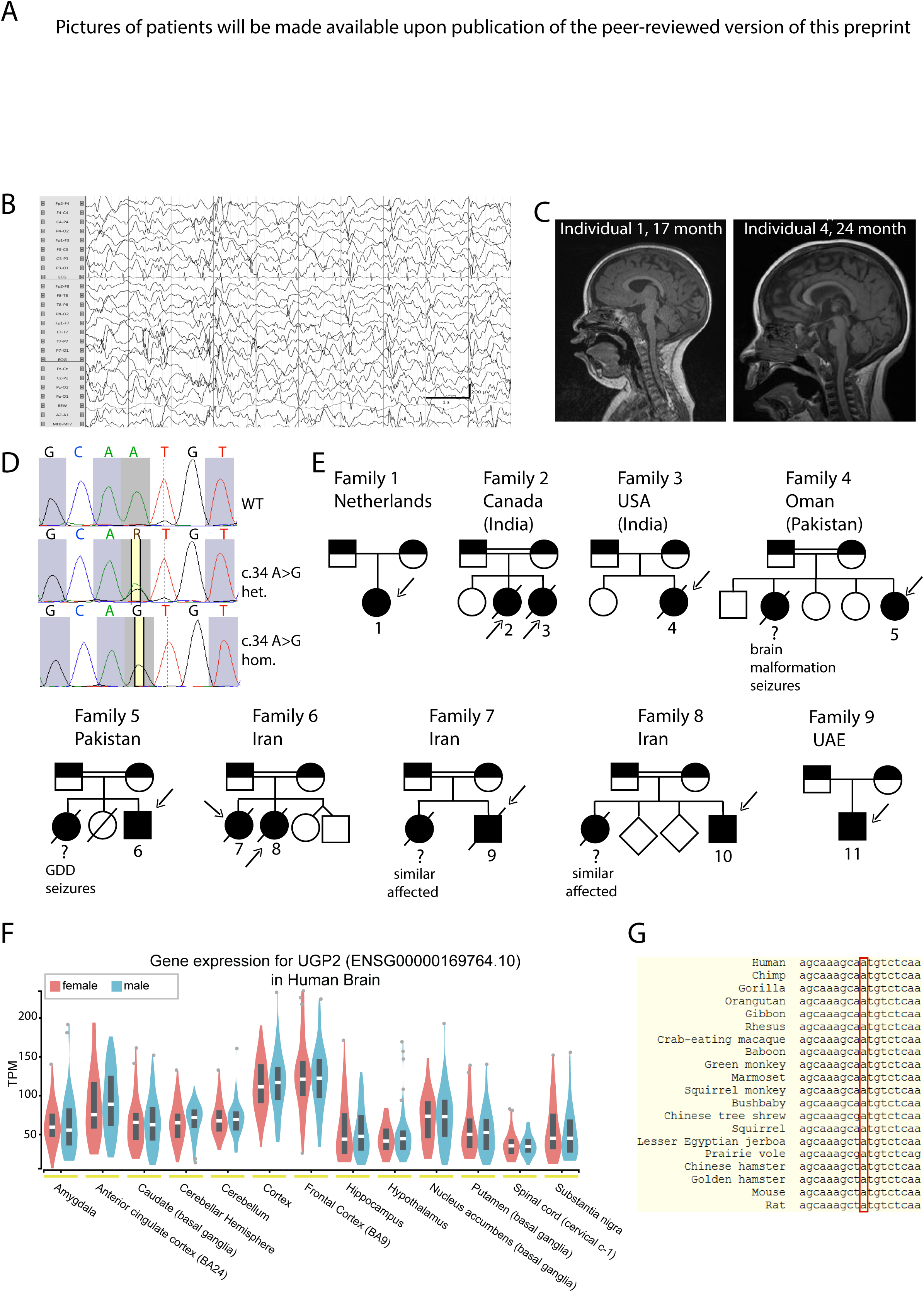
UGP2 homozygous variants in 10 individuals with severe epileptic encephalopathy A) Facial pictures of individual 1 (at 3, 18 and 23 month), individual 5 (at 9 years), individual 6 (at 11 month) and individual 10 (at 2 years). Note the progressive microcephaly with sloping forehead, suture ridging, bitemporal narrowing, high hairline, arched eyebrows, pronounced philtrum, a relatively small mouth and large ears. B) Electroencephalogram of individual 1 at the age of 8 month showing a highly disorganized pattern with high voltage irregular slow waves intermixed with multifocal spikes and polyspikes. C) T1-weighted mid sagittal brain MRI of individual 1 (age 17 month) and individual 4 (age 24 month) illustrating global atrophy and microcephaly but no major structural anomalies. D) Sanger sequencing traces of family 1, confirming the chr2:64083454A>G variant in *UGP2* in a heterozygous and homozygous state in parents and affected individual 1, respectively. E) Family pedigrees of ascertained patients. Affected individuals and heterozygous parents are indicated in black and half black, respectively. Affected individuals with confirmed genotype are indicated with an arrow, and numbers. Other affected siblings presenting with similar phenotypes are indicated with a question mark. Consanguineous parents are indicated with a double connection line. Male are squares, females circles; unknown sex indicated with rotated squares; deceased individuals are marked with a line. F) Violin plots showing distribution of gene expression (in TPM) amongst male and female samples from the GTEx portal^43^ for various brain regions. Outliers are indicated by dots G) Multiple species sequence alignment from the UCSC browser, showing that the ATG start site is highly conserved.

Having identified at least 12 individuals with an almost identical clinical phenotype and an identical homozygous variant in the same gene, led us to pursue *UGP2* as a candidate gene for a new genetic form of DEE. *UGP2* is highly expressed in various brain regions (**Figure 1F**), and also widely expressed amongst other tissues, including liver and muscle according to the data from the GTEx portal^35^ (**Supplementary Figure 1D**). The (chr2:64083454A>G) variant is predicted to cause a missense variant (c.34A>G, p.Met12Val) in *UGP2* isoform 1 (NM_006759), and to cause a translation start loss (c.1A>G, p.?.) of *UGP2* isoform 2 (NM_001001521), referred to as long and short isoform, respectively. The variant has not been reported in the Epi25 web browser^36^, ClinVar^37^, LOVD^38^, Exome Variant Server^39^, DECIPHER^40^, GENESIS^41^, GME variome^42^ or Iranome databases^43^, is absent from our in-house data bases and is found only 15 times in a heterozygous, but not homozygous, state in the 280,902 alleles present in *gnomAD* (MAF: 0.00005340)^44^. In the *GeneDx* unaffected adult cohort, the variant was found heterozygous 10 times out of 173,502 alleles (MAF: 0.00005764), in the ∼10,000 exomes of the Queen Square Genomic Center database two heterozygous individuals were identified, and out of 45,921 individuals in the *Centogene* cohort, 10 individuals are heterozygous for this variant. The identified variant has a CADD score (v1.4) of 19.22^45^ and Mutation Taster^46^ predicted this variant as disease causing. The nucleotide is strongly conserved over multiple species (**Figure 1G**). Analysis of WES data from 6 patients did provide evidence of a shared ROH between patients from different families, indicating that this same variant might represent an ancient mutation that originated some 26 generations ago (**Supplementary Figure 1C**). Interestingly, since most families originally came from regions of India, Pakistan and Iran, overlapping with an area called Balochistan, this could indicate that the mutation has originated there around 600 years ago. As Dutch traders settled in that area in the 17^th^ century, it is tempting to speculate that this could explain the co-occurrence of the variant in these distant places^47^.

### Short UGP2 isoform is predominantly expressed in brain and absent in patients with ATG mutations

Both UGP2 isoforms only differ by 11 amino acids at the N-terminal (**Figure 2A**) and are expected to be functionally equivalent^8^. To investigate how the A>G variant may cause DEE, we first obtained fibroblasts from individual 1 (homozygous for the A>G variant) and her heterozygous parents and analyzed the isoform expression by Western blotting (**Figure 2B**). Whereas the two isoforms were equally expressed in wild type fibroblasts, the expression of the shorter isoform was diminished to ∼25% of total UGP2 in heterozygous parents, both of individual 1 (**Figure 2B, C**) and of individual 2 and 3 (**Supplementary figure 2A, B**), and was absent in cells from the affected individual 1 (**Figure 2 B, C**; fibroblasts of the affected children in family 2 or other families were not available). Total UGP2 levels were not significantly different between the affected child and her parents, or between parents and wild type controls (**Figure 2D, Supplementary Figure 2C**). This indicates that the long isoform harboring the Met12Val missense variant is upregulated in fibroblast when the short isoform is missing. Moreover, this indicates that Met12Val does not affect the stability of the long isoform at the protein or transcript level (**Supplementary Figure 2D, E, F**). RNA-seq on peripheral blood samples of family 1 did not identify altered splicing events of *UGP2* and the global transcriptome of the proband was not different from her parents, although only a limited analysis could be performed as only a single sample was available for each individual (**Supplementary Figure 2G, H**). Both homozygous and heterozygous fibroblasts had a similar proliferation rate compared to wild type fibroblasts (**Figure 2E, Supplementary Figure 2I**), and immunocytochemistry confirmed a similar subcellular localization of UGP2 in mutant and wild type cells (**Figure 2F**). We then measured the enzymatic activity of UGP2 in wild type, heterozygous and homozygous fibroblasts, and found that mutant fibroblast had a similar capacity to produce UDP-glucose in the presence of exogenously supplied glucose-1-phosphate and UTP (**Figure 2G**). Altogether, this indicates that the long UGP2 isoform harboring the Met12Val missense change is functional and is therefore unlikely to contribute to the patient phenotype.

**Figure 2:**
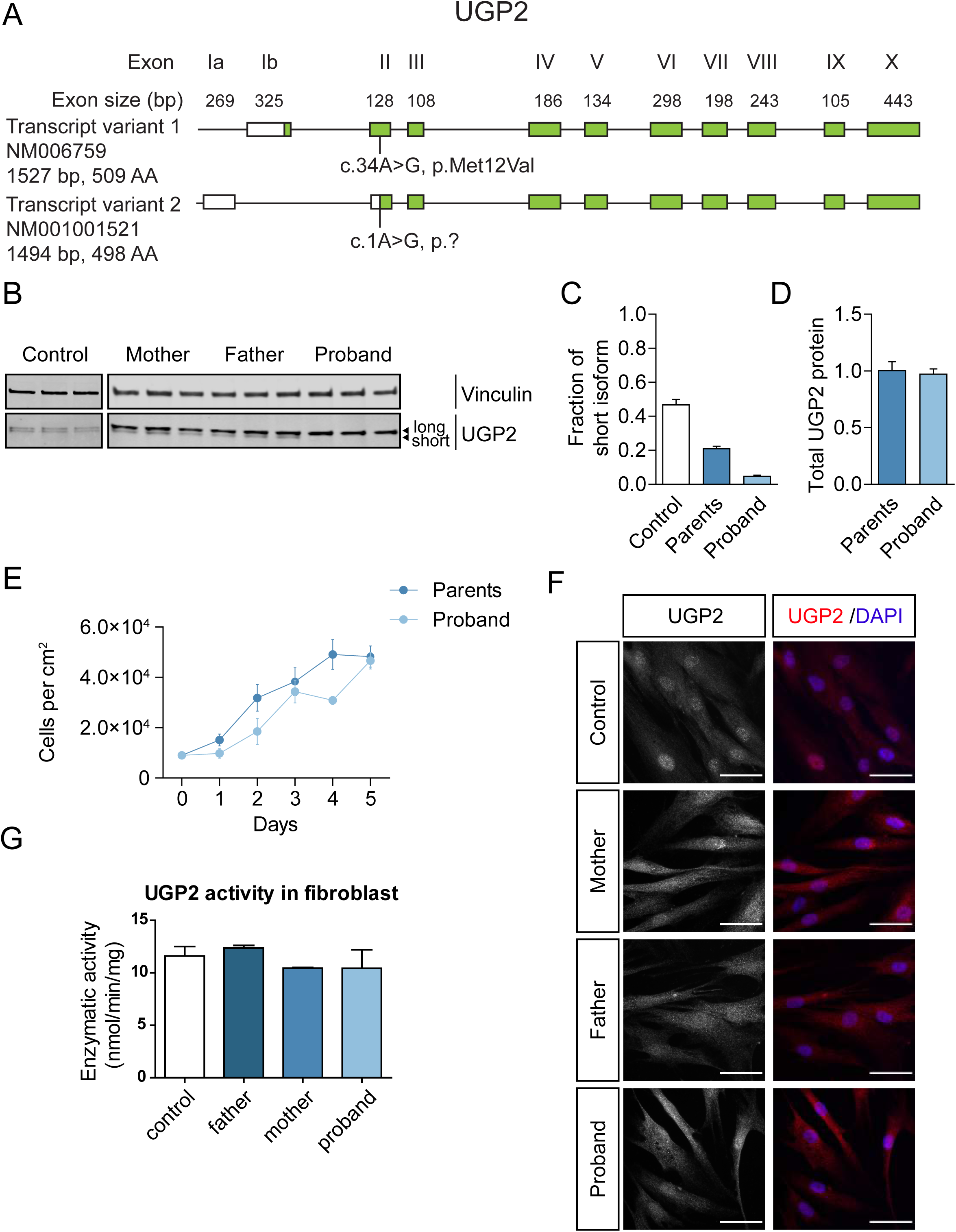
UGP2 homozygous variant leads to a loss of the shorter protein isoform in patient fibroblasts A) Schematic drawing of the human *UGP2* locus, with both long and short transcript isoforms. Boxes represent exons, with coding sequences indicated in green. The location of the recurrent mutation is indicated in both transcripts. B) Western blotting of cellular extracts derived from control fibroblasts or fibroblasts obtained from family 1, detecting the housekeeping control vinculin or UGP2. Note the two separated isoforms of UGP2 that have a similar intensity in wild type cells. The shorter isoform is less expressed in fibroblasts from heterozygous parents and absent in fibroblasts from the affected proband. C) Western blot quantification of the fraction of the short UGP2 protein isoform compared to total UGP2 expression in control, parental heterozygous and proband homozygous fibroblasts, as determined in three independent experiments. Error bars represent SEM. D) Western blot quantification of total UGP2 protein levels, as determined by the relative expression to the housekeeping control vinculin. Bar plot showing the results from three independent experiments. Error bars represent SEM; no significant differences were found between parents and proband, t-test, two-tailed. E) Cell proliferation experiment of fibroblast from heterozygous parents and homozygous proband from family 1, during a 5 days period, determined in three independent experiments. Error bars represent SEM. F) Immunocytochemistry on cultured control and UGP2 heterozygous and homozygous mutant fibroblast derived from family 1, detecting UGP2 (red). Nuclei are stained with DAPI. Scale bar = 50 µm. G) Enzymatic activity of UGP2 as measured in control and UGP2 heterozygous and homozygous mutant fibroblast derived from family 1. Shown is the mean of two independent experiments. Error bars represent SEM; no significant differences were found, unpaired t-test, two-tailed.

As the A>G variant results in a functional long UGP2 isoform but abolishes the translation of the shorter UGP2 isoform, we next investigated whether the ratio between short and long isoform differs amongst tissues. If so, the homozygous A>G variant would lead to depletion of UGP2 in tissues where mainly the short isoform is expressed, possibly below a threshold that is required for normal development or function. Western blotting on cellular extracts derived from wild type H9 human embryonic stem cells (ESCs), commercially acquired H9-derived neural stem cells (NSCs) and fibroblasts (**Figure 3A**) showed that, whereas the ratio between short and long isoform in fibroblasts was around 0.5, in ESCs it was 0.14 and in NSCs 0.77, indicating that the shorter UGP2 isoform is the predominant one in NSCs (**Figure 3B**). A similar trend was observed when assessing the transcript level, both by multiplex RT-PCR and RT-qPCR, using primers detecting specifically the short and long transcript isoform (**Figure 3C, D, E**). This indicates that differential isoform expression between cell types is regulated at the transcriptional level, possibly hinting at tissue-specific regulatory elements driving isoform expression. We next analyzed RNA-seq data from human fetal tissues^48-51^ to determine the fraction of reads covering short versus total *UGP2* transcripts (**Figure 3F**). This showed that in human fetal brain the short transcript isoform is predominantly expressed. To gain more insight into the cell type-specific expression of UGP2, we performed immunohistochemistry on human fetal brain tissues from the first to third trimester of pregnancy (**Figure 3G**). In the first trimester we found pale labeling of neuropil in the proliferative neuroepithelium of the hypothalamic, cortical, mesencephalic and thalamic regions (**Figure 3G-A/I, II, III, IV**), as well as the marginal zone of the spinal cord (**Figure 3G-A/V**) and cuboidal epithelial cells of choroid plexus (**Figure 3G-A/VI**). During the second trimester, UGP2 positivity was detected in neurons from the subplate region of the cerebral cortex (**Figure 3G-B/I, II**) and still in some of the cells in the neuroepithelium and subventricular zone (**Figure 3G-B/III**). Almost the same pattern of UGP2 distribution was found in the cerebral cortex of fetuses from the 3^rd^ trimester. Also, we found clear cytoplasmatic UGP2 expression in neurons from mesencephalic, inferior olivary and cerebellar nuclei during the second (**Figure 3G-B/IV, V, and VI**) and third trimester, respectively (**Figure 3G-C/IV, V**). In the white matter of the cerebellum in the third trimester, we identified single positive glial cells (**Figure 3G-C/VI**). In the cerebellar cortex we did not find specific positivity of cells on UGP2 (**Figure 3G-B, C/VII**). Cuboidal epithelial cells of choroid plexus preserved UGP2 positivity during the second trimester (**Figure 3G-B/VIII**) but lost it in the third trimester (**Figure 3G-C/VIII**). Together this indicates that UGP2 can be detected in a broad variety of cell types during brain development. On Western blotting, we noticed preferential expression of the shorter UGP2 isoform in the developing cortex and cerebellum from gestational weeks 14, 20 and 28 (**Figure 3H**) and in the frontal cortex of brains from weeks 21 and 23 (**Supplementary Figure 2J**). Together, this supports the hypothesis that the DEE phenotype in patients is caused by a major loss of functional UGP2 in the brain, as the short isoform represents virtually all UGP2 produced in this tissue.

**Figure 3:**
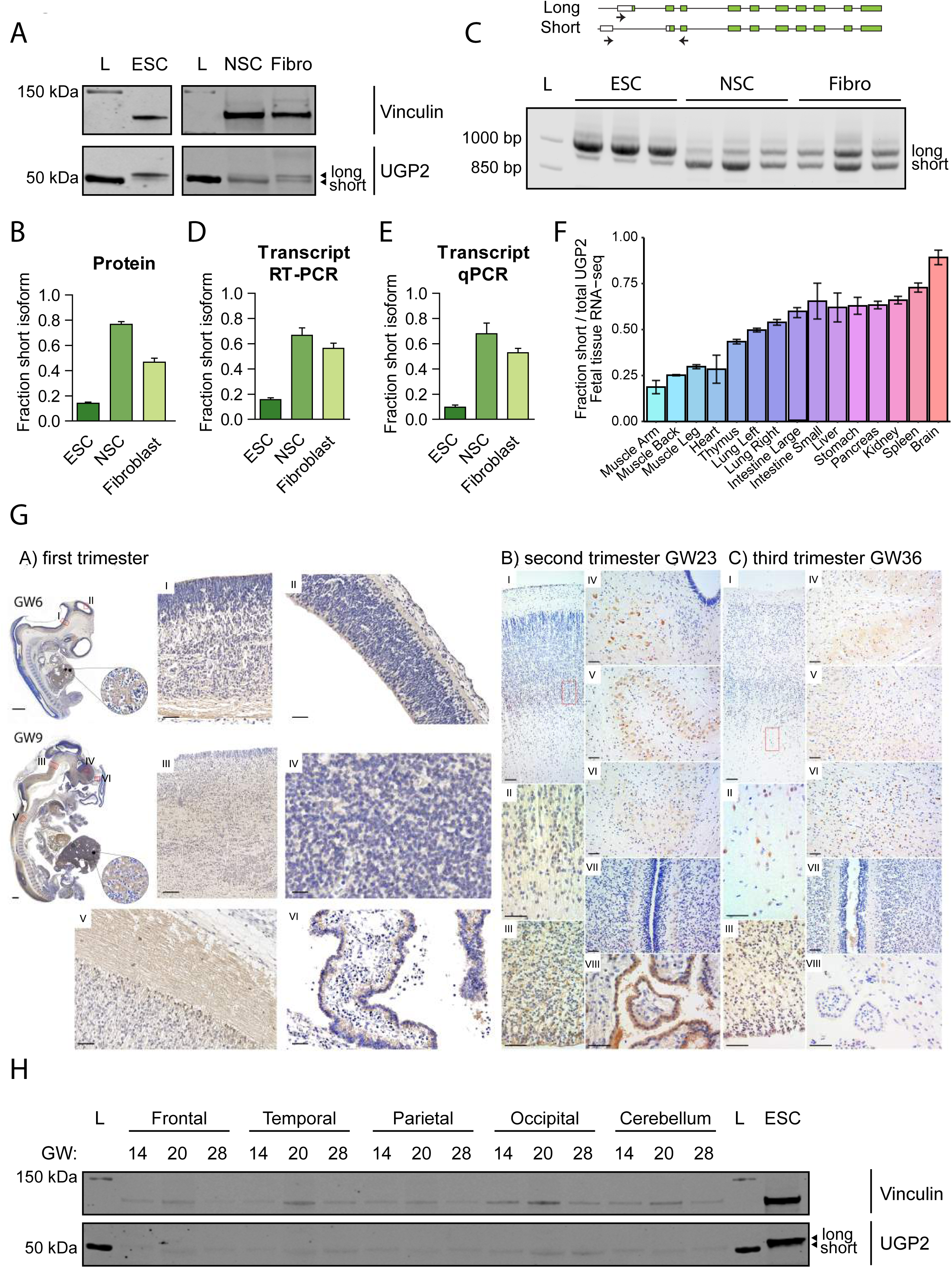
UGP2 short isoform is predominant in brain related cell types A) Western blotting showing UGP2 expression in H9 human embryonic stem cells (ESCs), H9 derived neural stem cells (NSCs) and fibroblasts (Fibro). Vinculin is used as a housekeeping control. Note the changes in relative expression between the two UGP2 isoforms in the different cell types. L, ladder. B) Western blot quantification of the fraction of the short UGP2 protein isoform compared to total UGP2 expression, as determined in three independent experiments. Error bars represent SEM. C) Multiplex RT-PCR of ESCs, NSCs and fibroblasts, showing a similar variability in isoform expression at the transcript as at the protein level. Each cell line was tested in triplicates. D) Quantification of the fraction of the short *UGP2* transcript isoform compared to total *UGP2* expression, from the multiplex RT-PCR from C). Error bars represent SEM E) Quantification of the fraction of the short *UGP2* transcript isoform compared to total *UGP2* expression by qRT-PCR in three independent experiments. Error bars represent SEM. F) Ratio of RNA-seq reads covering the short transcript isoform compared to the total reads (covering both short and long isoforms), in multiple fetal tissues. In RNA-seq samples derived from brain, virtually all UGP2 expression comes from the short isoform. Error bars represent SD. G) Immunohistochemistry detecting UGP2 in human fetal brains from the first, second and third trimester (gestational week (GW) 6, 9, 23 and 36). See text for details. H) Western blotting detecting UGP2 in various human brain regions at week 14, 20 and 28 of gestation, showing the virtual absence of the long isoform expression in fetal brain. Vinculin is used as a housekeeping control. L, ladder.

### Lack of the short UGP2 isoform leads to transcriptome changes upon differentiation into neural stem cells

To model the disease *in vitro*, we first engineered the homozygous A>G mutation in H9 ESCs to study the mutation in a patient independent genetic background and compare it to isogenic parental cells. We obtained two independent clones harboring the homozygous A>G change (referred to as knock-in, KI, mutant) and two cell lines harboring an insertion of an additional A after nucleotide position 42 of *UGP2* transcript 1 (chr2:64083462_64083463insA) (**Supplementary Figure 3A, B**) (referred to as knockout, KO). This causes a premature stop codon at amino acid position 47 (D15Rfs*33), leading to nonsense mediated mRNA decay and complete absence of UGP2 protein (**Supplementary Figure 3C**). All derived ESCs had a normal morphology and remained pluripotent as assessed by marker expression (**Supplementary Figure 3D, E**), indicating that the absence of UGP2 in ESCs is tolerated, in agreement with genome-wide LoF CRISPR screens which did not identify *UGP2* as an essential gene in ESCs^52,53^. We differentiated wild type, KI and KO ESCs into NSCs, using dual SMAD inhibition (**Supplementary Figure 4 A-C**). Wild type cells could readily differentiate into NSCs, having a normal morphology and marker expression, whereas differentiation of KI and KO cells was more variable and not all differentiations resulted in viable, proliferating NSCs. KO cells could not be propagated for more than 5 passages under NSC culture conditions (data not shown), which could indicate that the total absence of UGP2 protein is not tolerated in NSCs. When assessed by Western blotting, total UGP2 protein levels were reduced in KI cells and depleted in KO cells compared to wild type (**Supplementary Figure 4D, E**).

Next, we performed RNA-seq of wild type, KI and KO ESCs and NSCs to assess how depletion of UGP2 upon NSCs differentiation would impact on the global transcriptome (**Figure 4, Supplementary Figure 5, Supplementary Table 2**). In agreement with normal proliferation and morphology of KI and KO ESCs, all ESCs shared a similar expression profile of pluripotency associated genes and only few genes were differentially expressed between the three genotypes (**Supplementary Figure 5C, Supplementary Table 3**). This indicates that the absence of UGP2 in ESCs does not lead to major transcriptome alterations despite the central role of this enzyme in metabolism. Upon differentiation, cells from all genotypes expressed NSC markers (**Supplementary Figure 5F**), but when comparing wild type and KO cells, we observed noticeable changes, that were less pronounced in KI NSCs but still followed a similar trend (**Figure 4A, B, Supplementary Figure 5D, E**). Gene enrichment analysis showed that genes downregulated in KO and KI cells were implicated in processes related to the extra-cellular matrix, cell-cell interactions and metabolism, while genes upregulated in KO and KI cells were enriched for synaptic processes and genes implicated in epilepsy (**Figure 4C, Supplementary Table 4**). Both KO and KI cells showed an upregulation of neuronal expressed genes, indicating a tendency to differentiate prematurely. To validate RNA-seq findings, we tested several genes by RT-qPCR in wild type, KI and KO cells (**Figure 4D**). We also included KO rescue cells, in which we had restored the expression of either the wild type or the mutant UGP2 long isoform, leading each to an approximately 4-fold UGP2 overexpression at the NSC state compared to WT (**Supplementary Figure 4F**). Amongst the tested genes was *NNAT*, which showed a significant upregulation in KI and KO cells, which was rescued by restoration of UGP2 expression in KO NSCs. *NNAT* encodes neuronatin that stimulates glycogen synthesis by upregulating glycogen synthase and was previously found to be upregulated in Lafora disease. This lethal teen-age onset neurodegenerative disorder presenting with myoclonic epilepsy is caused by mutations in the ubiquitin ligase malin, leading to accumulation of altered polyglucosans^54^. Malin can ubiquitinate neuronatin leading to its degradation. As reduced UGP2 expression might impact on glycogen production, it seems plausible that this results in compensatory *NNAT* upregulation and in downstream aberrations contributing to the patient phenotypes. Indeed, neuronatin upregulation was shown to cause increased intracellular Ca^2+^ signaling, ER stress, proteasomal dysfunction and cell death in Lafora disease^55,56^, and was shown to be a stress responsive protein in the outer segment of retina photoreceptors^57,58^. Another interesting gene upregulated in KI and KO NSCs and downregulated in rescue cell lines was the autism candidate gene *FGFBP3*^59^. This secreted proteoglycan that enhances FGF signaling is broadly expressed in brain^60^, and functions as an extracellular chaperone for locally stored FGFs in the ECM, thereby influencing glucose metabolism by regulating rate-limiting enzymes in gluconeogenesis^61^. Other potentially relevant genes displaying the same expression trend were the heparan sulphate proteoglycan *GPC2* (a marker of immature neurons^62,63^), the helix-loop-helix transcription factor *ID4* (a marker of postmitotic neurons^64^), and the signaling molecule *FGFR3* that has been implicated in epilepsy^65^. Genes downregulated in KO cells and upregulated in rescue cells included urokinase-type plasminogen activator *PLAU* (deficiency in mouse models increases seizure susceptibility^66^), the glycoprotein *GALNT7* (upregulation of which has been found to promote glioma cell invasion^67^) and the brain tumor gene *MYBL1* (that has been shown to be regulated by *O*-linked *N*-acetylglucosamine^68^. Similar expression changes were observed in NSCs differentiated from induced pluripotent stem cells (iPSCs) that we had generated from family 1 (**Supplementary Figure 6)**. Together, RNA-seq showed that whereas the absence of UGP2 is tolerated in ESCs, its complete absence or reduced expression results in global transcriptome changes in NSCs, with many affected genes implicated in DEE relevant pathways.

**Figure 4:**
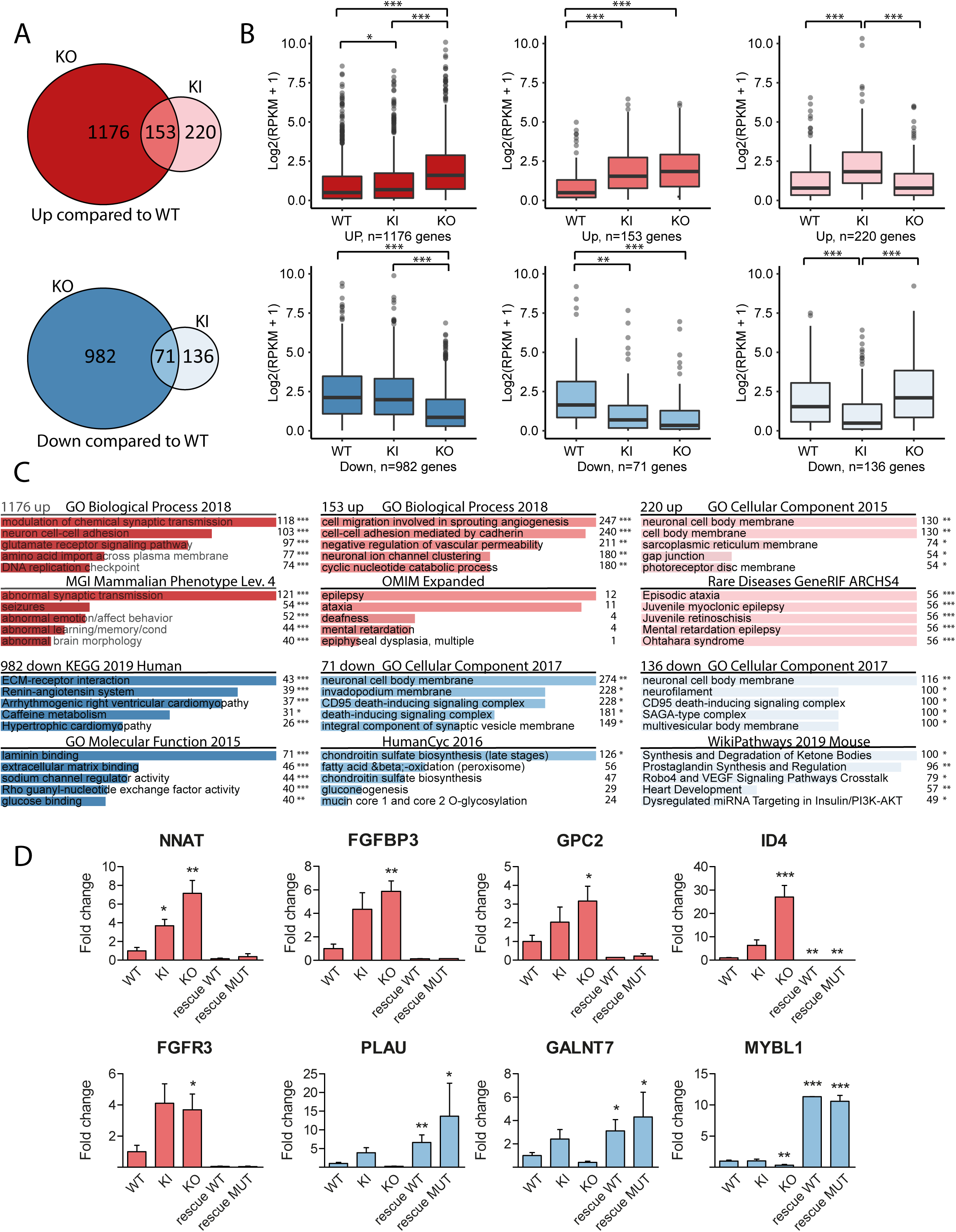
RNA-seq of UGP2 mutant H9 derived cell lines A) Venn diagram showing the overlap between differentially expressed genes in UGP2 KO or KI NSCs that are upregulated (upper panel genes with FDR<0.05 and LogFC>1) or downregulated (lower panel, genes with FDR<0.05 and LogFC<-1) compared to wild type NSCs. B) Box plot showing the distribution of gene expression levels (in Log2(RPKM+1)) from RNA-seq for the groups of genes displayed in **A**), in wild type, UGP2 KI or KO NSCs. Boxes are IQR; line is median; and whiskers extend to 1.5x the IQR (*=p<0.05; **=p<0.01,***=p<0.001, unpaired t-test, two-tailed). C) Enrichment analysis using Enrichr^119^ of up- or downregulated genes in NSCs from **A)** for selected gene ontology sets, showing the 5 most enriched terms per set. Combined score and p-value calculated by Enrichr are depicted (*=p<0.05; **=p<0.01; ***=p<0.001). D) qRT-PCR validation of differentially expressed genes from RNA-seq in wild type, UGP2 KI, UGP2 KO NSCs and KO NSCs that were rescued with either WT or MUT (Met12Val) transcript isoform 1, at p5 of NSC differentiation. Bar plot showing the mean fold change for the indicated genes compared to wild type, normalized for the housekeeping gene *TBP*. Results of two biological and two independent technical replicates are plotted. Colors match the Venn diagram group to which the tested genes belong to from A). Error bars represent SEM; (*=p<0.05; **=p<0.01,***=p<0.001, unpaired t-test, one-tailed).

### Absence of short UGP2 isoform leads to metabolic defects in neural stem cells

To investigate how reduced UGP2 expression levels in KO and KI cells would impact on NSC metabolism, we investigated the capacity to produce UDP-glucose in the presence of exogenously supplied glucose-1-phospate and UTP. KO NSCs showed a severely reduced ability to produce UDP-glucose (**Figure 5A)**. This reduction was rescued by ectopic overexpression of both long wild type and long mutant UGP2. KI cells showed a slightly reduced activity in ESCs (**Supplementary Figure 7A**), but a more strongly reduced activity in NSCs compared to wild type (**Figure 5A**), correlating with total UGP2 expression levels (**Supplementary Figure 4D, E**). Surprisingly, contrary to KO NSCs, KO ESC showed some residual capacity to produce UDP-glucose despite the complete absence of UGP2 (**Supplementary Figure 7A**). This could indicate that a yet to be identified enzyme can partially take over the function of UGP2 in ESCs but not NSCs, which might explain the lack of expression changes in this cell type upon UGP2 loss. iPSCs showed similar results (**Supplementary Figure 7B**). We next assessed the capacity to synthesize glycogen under low oxygen conditions by PAS staining, as it was previously shown that hypoxia triggers increased glycogen synthesis^69^. As expected, wild type ESCs cultured for 48 hours under hypoxia showed an intense cytoplasmic PAS staining in most cells (**Supplementary Figure 7C, D**), while KO ESCs showed a severely reduced staining intensity. This indicates that under hypoxia conditions, the residual capacity of ESC to produce UDP-glucose in the absence of UGP2 is insufficient to produce glycogen. KI ESCs were indistinguishable from wild type (**Supplementary Figure 7D)**. At the NSC state, many KO cells kept at low oxygen conditions for 48 hours died (data not shown) and those KO cells that did survive were completely depleted from glycogen granules (**Figure 5B, C**). This could be rescued by overexpression of both wild type or mutant long UGP2 isoform. KI NSCs showed a more severe reduction in PAS staining compared to the ESC state (**Figure 5B, C**), and we observed similar findings in patient iPSC derived NSCs (**Supplementary Figure 7E**). Together, this further indicates that upon neural differentiation the isoform expression switch renders patient cells depleted of UGP2, leading to a reduced capacity to synthesize glycogen. This can directly be involved in the DEE phenotype, as, besides affecting energy metabolism, reduction of glycogen in brain has been shown to result in I) impairment of synaptic plasticity^70^; II) reduced clearance of extracellular potassium ions leading to neuronal hypersynchronization and seizures^71-73^; and III) altered glutamate metabolism^74^. To investigate how reduced UDP-glucose levels would impact on glycosylation, we next, investigated glycosylation levels by means of LAMP2, a lysosomal protein known to be extensively glycosylated both by N-linked and O-linked glycosylation^75^. We found that KO NSCs show hypoglycosylation of LAMP2 that is rescued by the over expression of both WT and mutant long isoform (**Figure 5D**). In contrast, in ESCs no glycosylation defects were noticed (**Supplementary Figure 7F**). Finally, we investigated whether the absence of UGP2, affecting protein glycosylation, could induce ER stress and thus unfolded protein response (UPR). Whereas in ESCs, the absence of UGP2 did not result in a detectable effect on UPR markers (**Supplementary Figure 7G**), in NSCs we noticed an increased expression of these genes both in KO and in KI cells (**Figure 5E**). This indicates that NSCs having UGP2 levels under a certain threshold are more prone to ER-stress and UPR. In agreement with this, we did not observe upregulation of UPR markers in patient derived fibroblast, which have similar total UGP2 expression levels compared to controls (**Supplementary Figure 7H**). Together this indicates that upon differentiation to NSCs, KI cells become sufficiently depleted of UGP2 to have reduced synthesis of UDP-glucose, leading to defects in glycogen synthesis and protein glycosylation and to the activation of UPR response. Alterations of these crucial processes are likely to be implicated in the pathogenesis leading to increased seizure susceptibility, altered brain microstructure and progressive microcephaly.

**Figure 5:**
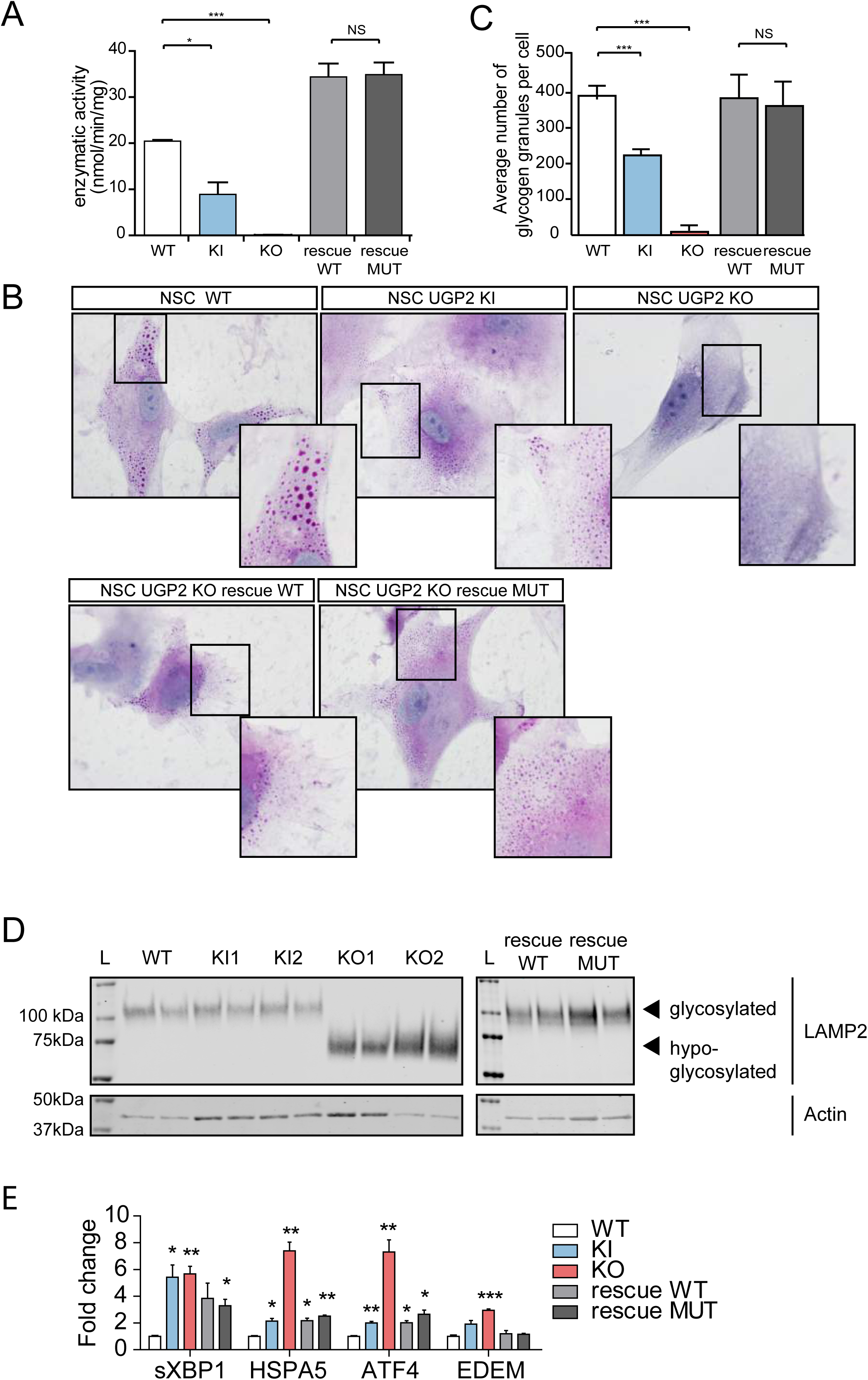
metabolic changes upon UGP2 loss A) UGP2 enzymatic activity in WT, UGP2 KI, KO and KO NSCs rescued with wildtype or mutant Met12Val isoform 1 of UGP2. Bar plot showing the mean of two replicate experiments, error bar is SEM. *=p<0.05; ***=p<0.001, unpaired t-test, two-tailed. B) Representative pictures of PAS staining in WT, KI, KO and rescue NSCs. Nuclei are counterstained with hematoxylin (blue). Inserts show zoom-in of part of the cytoplasm. Note the presence of glycogen granules in WT NSCs, their diminished number in KI NSCs, their absence in KO NSCs and their reappearance upon rescue with both wild type long UGP2 as with Met12Val long UGP2. C) Quantification of the number of glycogen granules per cell in WT, UGP2 KI, KO and rescue NSCs, after 48 hours culture under low-oxygen conditions. Shown is the average number of glycogen granules per cell, n=80-100 cells per genotype. Error bars represent the SD. ***=p<0.001, unpaired t-test, two-tailed. D) Western blotting detecting LAMP2 (upper panel) and the housekeeping control ACTIN (lower panel) in cellular extracts from ESC-derived NSCs, that are wt, UGP2 KI, KO and KO cells rescued with either the long wildtype isoform 1 or the mutant Met12Val isoform 1. Glycosylated LAMP2 runs at ∼110 kDa, whereas hypo-glycosylated LAMP2 is detected around 75 kDa. The absence of detectable changes in LAMP2 glycosylation in KI cells is likely explained by a non-complete isoform switch upon in vitro NSC differentiation, resulting in residual UGP2 levels (c.p**. Supplementary Figure 5D**). E) qRT-PCR expression analysis for UPR marker genes (spliced *XBP1, HSPA5, ATF4* and *EDEM*) in WT, KI, KO and rescue NSCs. Shown is the mean fold change for the indicated genes compared to wild type, normalized for the housekeeping gene *TBP*. Results of two biological and two independent technical replicates are plotted, from two experiments. Error bars represent SEM; *=p<0.05; **=p<0.01,***=p<0.001, unpaired t-test, two-tailed

### Ugp2a and Ugp2b double mutant zebrafish recapitulate metabolic changes during brain development, have an abnormal behavioral phenotype, visual disturbance, and increased seizure susceptibility

Finally, to model the consequences of the lack of UGP2 *in vivo*, we generated zebrafish mutants for both *ugp2a* and *ugp2b*, the zebrafish homologs of *UGP2*, using CRISPR-Cas9 injections in fertilized oocytes in a background of a radial glia/neural stem cell reporter^76^. Double homozygous mutant lines having frameshift deletions for both genes confirmed by Sanger sequencing could be generated but the only viable combination, obtained with *ugp2a* loss, created a novel ATG in exon 2 of ugp2b, leading to a hypomorphic allele (**Figure 6A**). Homozygous *ugp2a/b* mutant zebrafish had a normal gross morphology of brain and radial glial cells (**Figure 6B**), showed a largely diminished activity to produce UDP-glucose in the presence of exogenously supplied glucose-1-phospate and UTP (**Figure 6C**), and showed a reduction in c-FOS expression levels, indicating reduced global neuronal activity (**Figure 6D**). To monitor possible spontaneous seizures, we performed video tracking experiments of developing larvae under light-dark cycling conditions at 5 days post fertilization (dpf). Control larvae show increased locomotor activity under light conditions, and although *ugp2* double mutant larvae still responded to increasing light conditions, they showed a strongly reduced activity (**Figure 6E, F**). This could indicate that their capability to sense visual cues is diminished, or that their tectal processing of visual input is delayed, resulting in reduced movements. Strikingly, upon careful inspection, we noticed that *ugp2* double mutant larvae did not show spontaneous eye movements, in contrast to age-matched control larvae (**Figure 6G, Supplemental Movie 1 and 2**). Whereas we did not observe an obvious spontaneous epilepsy phenotype in these double mutant larvae, upon stimulation with 4-aminopyridine (4-AP), a potent convulsant, double mutant larvae showed an increased frequency and duration of movements at high velocity compared to controls, which might indicate an increased seizure susceptibility (**Figure 6H, I**). Taken together, severely reduced Ugp2a/Ugp2b levels result in a behavior defect with reduced eye movements, indicating that also in zebrafish Ugp2 plays an important role in brain function.

**Figure 6:**
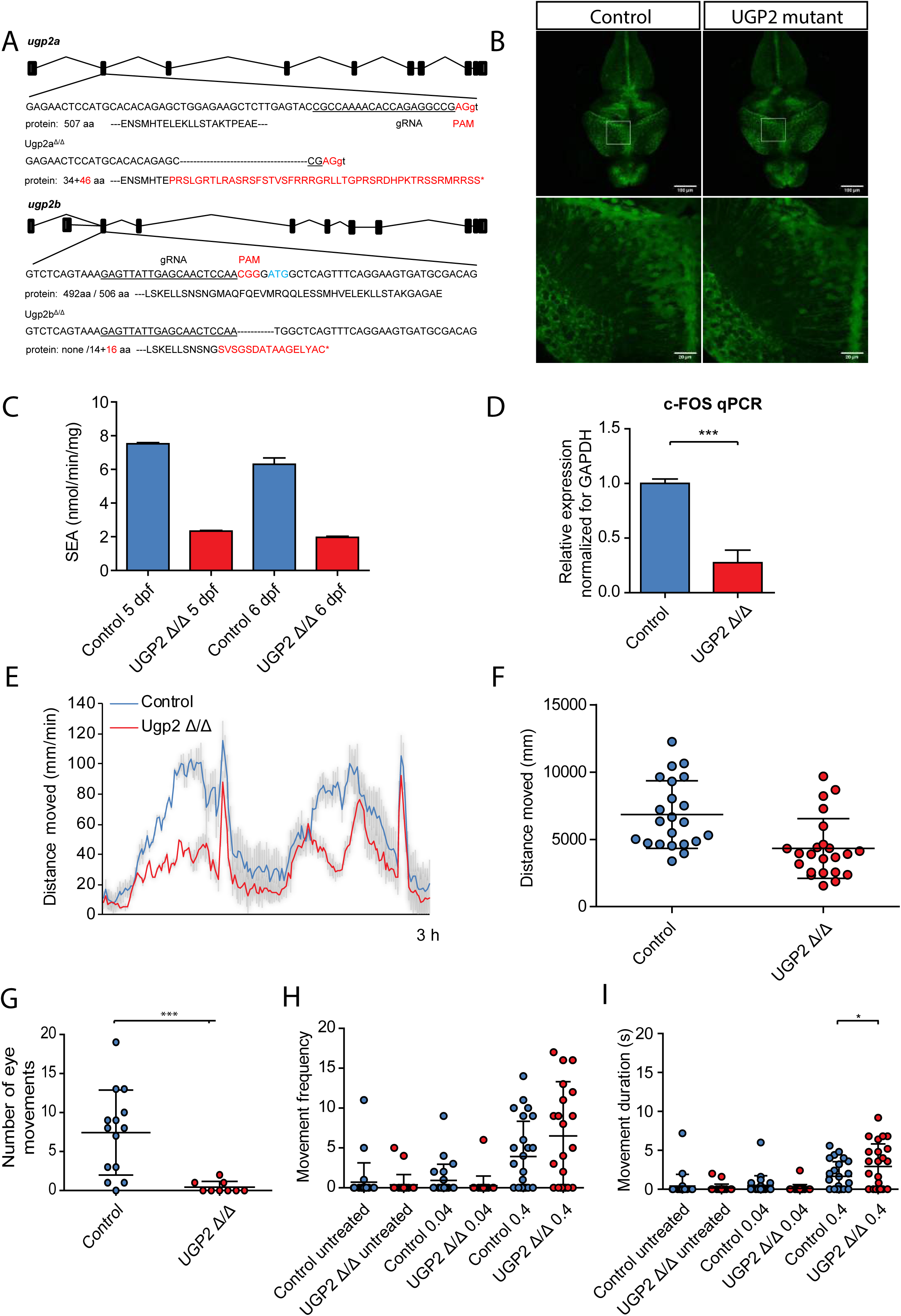
zebrafish disease modelling A) Schematic drawing of the *ugp2a* and *ugp2b* loci in zebrafish and the generated mutations indicated. B) Confocal images (Maximum projection of confocal Z-stacks) of the brain of wild type (left) and *ugp2a*^Δ/Δ^; *ugp2b*^Δ/Δ^ mutant zebrafish larvae (right), both in an *slc1a2b*-citrine reporter background, at 4 days post fertilization (dpf). IThe lower panels are higher magnifications of the boxed regions indicated in the upper panels. Scale bar in upper panel is 100 µm, in lower panel 20 µm. In upper panel, Z = 45 with step size 4 µm; In lower panel, Z = 30 with step size 2 µm. C) Enzymatic activity in *ugp2* double mutant zebrafish larvae at 4 and 5 dpf, compared to wild type age matched controls, showing reduced Ugp2 enzyme activity in double mutant zebrafish. D) qRT-PCR for the neuronal activity marker c-FOS in wild type and *ugp2* double mutant larvae at 3 dpf. For each group, 2 batches of 12 larvae were pooled. Shown is the mean fold change for the indicated genes compared to wild type, normalized for the housekeeping gene *gapdh*. Error bars represent SEM; ***= p<0.001, unpaired *t*-test, two-tailed. E) Representative graph of a locomotion assay showing the total distance moved by larvae during the dusk-dawn routine (total time: 3 hr 12 min), n = 24 larvae per genotype. Grey shading shows the standard error of the mean. F) Quantification of the total distance moved throughout the experiment from E) excluding the dark period. G) Quantification of the number of observed spontaneous eye movements during a 2 minutes observation in wild type and *ugp2* double mutant larvae at 4 dpf. Each dot represents one larva; shown is the average and SD; ***p<0.001, *t*-test, two tailed. H) Quantification of the frequency of movements at a speed of > 15 mm/s, for wild type control and *ugp2* double mutant zebrafish larvae at 4 dpf, treated with mock control or with 0.04 nM or 0.4 nM 4-AP during a 35 minutes observation. Each dot represents a single larva; results of two experiments are shown, with in total 24 larvae per condition. I) As H, but now assessing movement duration at a speed of > 15 mm/s. * =p<0.05, two way ANOVA with Bonferoni post-test.

### UGP2 is an essential gene in humans and ATG mutations of tissue specific isoforms of essential genes potentially cause more rare genetic diseases

Several lines of evidence argue that UGP2 is essential in humans. First, no homozygous LoF variants or homozygous exon-covering deletions for *UGP2* are present in *gnomAD or GeneDx controls*, and homozygous variants in this gene are limited to non-coding changes, synonymous variants and 5 missense variants, together occurring only 7 times homozygous (**Supplementary Table 5**). Also, no homozygous or compound heterozygous *UGP2* LoF variants were found in published studies on dispensable genes in human knockouts^77-79^, or in the *Centogene* (*CentoMD*) or *GeneDx* patient cohorts, encompassing together many thousands of individuals, further indicating that this gene is intolerant to loss-of-function in a bi-allelic state. In addition, no homozygous deletions of the region encompassing *UGP2* are present in DECIPHER^40^ or ClinVar^37^. Second, *UGP2* has been identified as an essential gene using gene-trap integrations^80^ and in CRISPR-Cas9 LoF screens in several human cell types^81-85^. Finally, studies in yeast^86,87^, fungus^88^ and plants^89-91^ consider the orthologs of *UGP2* as essential, and the absence of *Ugp2* in mice is predicted to be lethal^92^. In flies, homozygous UGP knock-outs are lethal while only hypomorphic compound heterozygous alleles are viable but have a severe movement defect with altered neuromuscular synaptogenesis due to glycosylation defects^93^. To further investigate the essentiality of UGP2, we performed differentiation experiments of our WT, KO and rescue ESCs. Differentiation of KO ESCs into hematopoietic stem cells (HSCs) resulted in severe downregulation of *GATA2* compared to wild type cells, and this was restored in rescue cell lines (**Figure 7A**). GATA2 is a key transcription factor in the developing blood system, and knockout of *Gata2* is embryonic lethal in mice due to defects in HSC generation and maintenance^94,95^. Differentiation of ESCs into cardiomyocytes similarly affected key marker gene expression in KO cells, and these changes were restored upon UGP2 rescue (**Figure 7B, C**). Whereas WT ESCs could generate beating cardiomyocytes after 10 days, these were not seen in KO ESCs. Taken together this argues that the complete absence of UGP2 in humans is probably incompatible with life, a hypothesis that cannot be tested directly. However, if true, this could well explain the occurrence of the unique recurrent mutation in all cases presented herein. Given the structure of the *UGP2* locus (**Figure 2A**), every LoF variant would affect either the long isoform, when located in the first 33 nucleotides of the cDNA sequence, or both the short and long isoform when downstream to the ATG of the short isoform. Therefore, the short isoform start codon is the only mutational target that can disrupt specifically the short isoform. In this case, the Met12Val change introduced into the long isoform does not seem to disrupt UGP2 function to such an extent that this is intolerable and therefore allows development to proceed for most tissues. However, the lack of the short UGP2 isoform caused by the start codon mutation results in a depletion of functional UGP2 in tissues where normally the short isoform is predominantly expressed. In brain this reduction diminishes total UGP2 levels below a threshold for normal development, causing a severe epileptic encephalopathy syndrome. Given the complexity of the human genome with 42976 transcripts with RefSeq peptide IDs, perhaps also other genetic disorders might be caused by such tissue restricted depletion of essential proteins. Using a computational homology search of human proteins encoded by different isoforms, we have identified 1766 genes that share a similar structure to the *UGP2* locus (e.g. a shorter protein isoform that is largely identical to the longer protein isoform, translated from an ATG that is contained within the coding sequence of the long isoform) (**Figure 7D**). When filtering these genes for 1) those previously shown to be essential^6^, 2) not associated with disease (e.g. no OMIM phenotype) and 3) those proteins where the shorter isoform is no more than 50 amino acids truncated at the N-terminal compared to the longer isoform, we identified 247 genes (**Supplementary Table 6**). When comparing the ratios of isoform specific reads obtained from different fetal RNA-seq data^48-51^ we noticed that many of these genes show differential isoform expression amongst multiple tissues, with many genes showing either expression of the long or the short isoform in a particular tissue (**Figure 7E**). Homozygous LoF variants or start codon altering mutations in these genes are rare in *gnomAD* (**Supplementary Table 7**), and it is tempting to speculate that mutations in start codons of these genes could be associated with human genetic diseases, as is the case for *UGP2*. Using mining of data from undiagnosed patients from our own exome data base, the Queen Square Genomic Center database and those from *Centogene* and *GeneDx*, we found evidence for several genes out of the 247 having rare, bi-allelic variants affecting the start codon of one of the isoforms that could be implicated in novel disorders (*unpublished observations*) and give one such example in the **Supplementary Note**. Together, these findings highlight the relevance of mutations resulting in tissue-specific protein loss of essential genes for genetic disorders.

**Figure 7:**
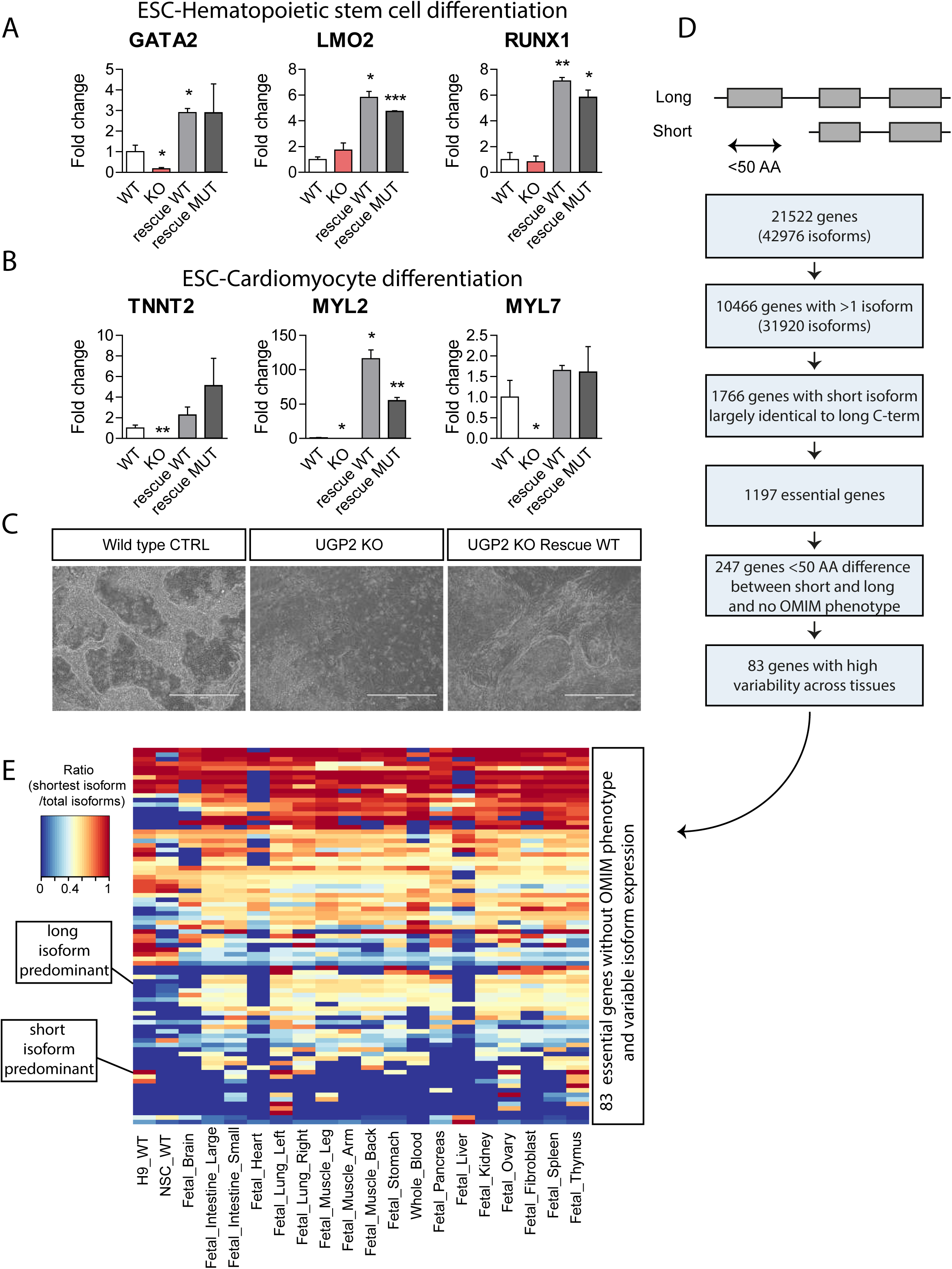
Essentiality of UGP2 and other disease candidate genes with a similar mutation mechanism A) qRT-PCR analysis of the hematopoietic stem cell markers *GATA2, LMO2* and *RUNX1*, after 12 days of differentiation of wild type, UGP2 KO and UGP2 KO rescue ESCs. Shown is the mean fold change for the indicated genes compared to wild type, normalized for the housekeeping gene *TBP*. Results of two biological and two technical replicates are plotted. Error bars represent SEM; *=p<0.05; **=p<0.01,***=p<0.001, unpaired t-test, two-tailed. B) As A), but now for cardiomyocyte differentiation at day 15, assessing expression of the cardiomyocyte markers *TNNT2, MYL2* and *MYL7*. C) Bright-field image of cardiomyocyte cultures of wild type, UGP2 KO and rescue cells. Note the elongated organized monolayer structures cardiomyocytes capable of beating in wild type and rescue cells, that are absent in KO cultures. Scale bar is 400 µm. D) Scheme showing the homology search to identify genes with a similar structure as UGP2, where ATG altering mutations could affect a tissue specific isoform causing genetic disease. E) Heat map showing the ratio of short isoform expression over total isoform expression from published RNA-seq data amongst 20 tissues for 83 out 247 essential genes that are not yet implicated in disease and in which the short and longer protein isoform differ by less than 50 amino acids at the N-terminal.

## Discussion

Here we describe a recurrent variant in 12 individuals from 10 families, affecting the start codon of the shorter isoform of the essential gene *UGP2* as a novel cause of a severe DEE. Using *in vitro* and *in vivo* disease modeling, we provide evidence that the reduction of *UGP2* expression in brain cells leads to global transcriptome changes, a reduced ability to produce glycogen, alterations in glycosylation and increased sensitivity to ER stress, which together can explain the phenotype observed in the patients. Most likely our findings *in vitro* underestimate the downstream effects in patient cells, as in fetal brain the longer isoform expression is almost completely silenced and virtually all UGP2 comes from the shorter isoform, which in patient cells cannot be translated. During our *in vitro* NSC differentiation this isoform switch is less complete, leaving cells with the patient mutation with some residual UGP2. Strikingly, the clinical phenotype seems to be very similar in all cases, including intractable seizures, absence of developmental milestones, progressive microcephaly and a disturbance of vision, with retinal pigment changes observed in all patients who had undergone ophthalmological examination. Also, all patients seem to share similar, although mild, dysmorphisms, possibly making this condition a recognizable syndrome.

The involvement of *UGP2* in genetic disease is surprising. Given its central role in nucleotide-sugar metabolism it is expected that loss of this essential protein would be incompatible with life, and therefore loss-of-function should not be found in association with postnatal disease. Our data argue that indeed a total absence of UGP2 in all cells is lethal, but that tissue-specific loss, as caused here by the start codon alteration of an isoform important for brain, can be compatible with postnatal development but still results in a severe phenotype. Given that any other LoF variant across this gene would most likely affect both protein isoforms, this could also explain why only a single mutation is found in all individuals. The fact that the Met12Val long isoform was able to rescue the full KO phenotype indicates that the missense change introduced to the long protein isoform does not affect UGP2 function. As other variants at this start codon, even heterozygous, are not found, possibly missense variants encoding for leucine, lysine, threonine, arginine or isoleucine (e.g. amino acids that would be encoded by alternative changes affecting the ATG codon) at this amino acid location in the long isoform could not produce a functional protein and are therefore not tolerated. Although start codon mutations have previously been implicated in disease^96,97^, there are no reports, to our knowledge, on disorders describing start codon alterations of other essential genes, leading to alterations of tissue specific isoforms. Using a genome-wide homology search, we have identified a large list of other essential genes with a similar locus structure and variable isoform expression amongst tissues, where similar ATG altering variants could affect tissue-relevant expression. An intriguing question is why evolution has resulted in a large number of genes encoding almost identical protein isoforms. It will be interesting to further explore the mutational landscape of these genes in cohorts of currently unexplained patients.

## Author contributions

EP performed molecular biology experiments, with help from AN and DP. HvdL, WB and TvH performed zebrafish work. PvdB and EHJ performed enzymatic analyses. IC performed brain immunohistochemistry and supplied tissues samples. EA supplied tissue samples. MG generated iPSCs. WvI and WGdV performed and SY analysed RNA-seq. SY performed gene homology search. Patient recruitment and diagnosing was performed in the different families as follows: Family 1: TSB, ASB, and EM phenotyped patient 1, MvS analysed WES; Family 2: LB and MK phenotyped patient 2 and 3, KGM, AB, KR analyzed WES; Family 3: JNK and JB phenotyped patient 4, KGM, AB, KR analyzed WES. Family 4: AaF, FaM, RM and FaA phenotyped patient 5, EJK analyzed WES; Family 5: FZ and NR phenotyped patient 6, SE, HH analyzed WES; Family 6, family 7 and family 8: MM, AE, ZK, FMD, MD, EGK phenotyped patients 7-10, JV, RM, HH analyzed WES; Family 9: JH phenotyped patient 11, KKK, ABA analysed WES; RT, KR, KKK, PB, ABA, RM, HH provided genetic data for population analysis. TSB identified patient 1, conceived the study, obtained funding, supervised the lab work and wrote the manuscript, with input from all main authors. All authors approved the final version of the manuscript.

## Acknowledgements

We are indebted to the parents of the patients for their kind cooperation. We thank Virginie Verhoeven and Gerben Schaaf for critically reading our manuscript and Grazia Mancini for helpful discussions. We thank Gerben Schaaf for providing the LAMP2 antibody, and Eskeatnaf Mulugeta for bioinformatics advice. DP was supported by an Erasmus+ Traineeship Programme. HH is supported by the Rosetree Trust, Ataxia UK, MSA Trust, Brain Research UK, Muscular Dystrophy UK, Muscular Dystrophy Association (MDA USA), Higher Education Commission of Pakistan, The MRC (MR/S01165X/1, MR/S005021/1, G0601943), Wellcome Trust (WT093205MA, WT104033AIA and the Synaptopathies Strategic Award, 165908) and the National Institute for Health Research University College London Hospitals Biomedical Research Centre. Families 5, 6, 7 and 8 were collected as part of the SYNaPS Study Group collaboration funded by The Wellcome Trust and strategic award (Synaptopathies) funding. Research for these families was conducted as part of the Queen Square Genomics group at University College London, supported by the National Institute for Health Research University College London Hospitals Biomedical Research Centre. TVH is supported by an Erasmus University Rotterdam (EUR) fellowship. TSB’s lab is supported by the Netherlands Organisation for Scientific Research (ZonMW Veni, grant 91617021), a NARSAD Young Investigator Grant from the Brain & Behavior Research Foundation, an Erasmus MC Fellowship 2017 and Erasmus MC Human Disease Model Award 2018. TSB, IC and EA acknowledge support from COST action CA16118 that facilitated this collaboration.

## Online Methods

### Patient recruitment

All affected probands were investigated by their referring physicians and all genetic analysis was performed in a diagnostic setting. Legal guardians of affected probands gave informed consent for genomic investigations and publication of their anonymized data.

### Next generation sequencing of index patients

#### Individual 1

Genomic DNA was isolated from peripheral blood leukocytes of proband and both parents and exome-coding DNA was captured with the Agilent Sure Select Clinical Research Exome (CRE) kit (v2). Sequencing was performed on an Illumina HiSeq 4000 with 150bp paired end reads. Reads were aligned to hg19 using BWA (BWA-MEM v0.7.13) and variants were called using the GATK haplotype caller (v3.7 (reference: http://www.broadinstitute.org/gatk/)^98^. Detected variants were annotated, filtered and prioritized using the Bench lab NGS v5.0.2 platform (Agilent technologies). Initially, only genes known to be involved in epilepsy were analyzed, followed by a full exome analysis revealing the homozygous UGP2 variant

#### Individuals 2, 3 and 4

Using genomic DNA from the proband and parents (individual 4) or the proband, parents, and affected sibling (individual 2 and 3), the exonic regions and flanking splice junctions of the genome were captured using the SureSelect Human All Exon V4 (50 Mb) (individual 4) or the IDT xGen Exome Research Panel v1.0 (individual 2 and 3). Massively parallel (NextGen) sequencing was done on an Illumina system with 100bp or greater paired-end reads. Reads were aligned to human genome build GRCh37/UCSC hg19, and analyzed for sequence variants using a custom-developed analysis tool. Additional sequencing technology and variant interpretation protocol has been previously described^99^. The general assertion criteria for variant classification are publicly available on the GeneDx ClinVar submission page (http://www.ncbi.nlm.nih.gov/clinvar/submitters/26957/)

#### Individual 5 (Nijmegen)

Diagnostic exome sequencing was done at the Departments of Human Genetics of the Radboud University Medical Center Nijmegen, The Netherlands and performed essentially as described previously^100^.

#### Individual 6, 7, 8, 9 and 10

After informed consent, we collected blood samples from the probands, their parents and unaffected siblings, and extracted DNA using standard procedures. To investigate the genetic cause of the disease, WES was performed in the affected proband. Nextera Rapid Capture Enrichment kit (Illumina) was used according to the manufacturer instructions. Libraries were sequenced in an Illumina HiSeq3000 using a 100-bp paired-end reads protocol. Sequence alignment to the human reference genome (UCSC hg19), and variants calling, and annotation were performed as described elsewhere^101^. After removing all synonymous changes, we filtered single nucleotide variants (SNVs) and indels, only considering exonic and donor/acceptor splicing variants. In accordance with the pedigree and phenotype, priority was given to rare variants [<1% in public databases, including 1000 Genomes project, NHLBI Exome Variant Server, Complete Genomics 69, and Exome Aggregation Consortium (ExAC v0.2)] that were fitting a recessive or a de novo model.

#### Individual 11 and 12

Whole exome sequencing was performed at CENTOGENE AG, as previously described^102^.

### Human brain samples

Tissue was obtained, upon informed consent, and used in a manner compliant with the Declaration of Helsinki and the Research Code provided by the local ethical committees. Fetal brains were preserved after spontaneous or induced abortions with appropriate maternal written consent for brain autopsy and use of rest material for research. We performed a careful histological and immunohistochemical analysis and evaluation of clinical data (including genetic data, when available). We only included specimens displaying a normal cortical structure for the corresponding age and without any significant brain pathology.

### Brain tissue immunohistochemistry

For immunohistochemical analysis, we used 2 cases from the first trimester (GW6 and GW9), 4 cases from the second trimester (GW21, GW23, GW24 and GW26) and 2 cases from the third trimester (GW33 and GW36). Anatomical regions were determined according to the atlas of human brain development^103-106^. We cut 4 µm sections from formalin-fixed, paraffin embedded whole fetuses (GW6 and GW9) and brain tissue from cerebral, mesencephalic, cerebellar and brain stem region (from GW21 to GW36). Slides were stained with mouse anti-UGP2 (C-6) in a 1:150 dilution (Santa Cruz) and visualized using Mouse and Rabbit Specific HRP/DAB (ABC) Detection IHC kit (Abcam). Mayer’s hematoxylin was used as a counterstain for immunohistochemistry followed by mounting and coverslipping (Bio-Optica) for slides. Prepared slides were analyzed and scanned under a VisionTek^®^ Live Digital Microscope (Sakura).

### Cloning of UGP2 cDNA

RNA was isolated using TRI reagent (Sigma) from whole peripheral blood of index patient 1 and her parents, after red blood cell depletion with RBC lysis buffer (168mM NH_4_Cl, 10mM KHCO_3_, 0.1mM EDTA). cDNA was synthesized following the iSCRIPT cDNA Synthesis Kit (Bio-Rad) protocol, and the coding sequence of the long and short UGP2 isoform (wild type or mutant) was PCR-amplified together with homology arms for Gibson assembly (see **Supplementary Table 8** for primer sequences) using Phusion High-Fidelity DNA polymerase (NEB). PCR amplified DNA was then cloned by Gibson assembly as previously described^107^ in a pPyCAG-IRES-puro plasmid (a kind gift of Ian Chambers, Edinburgh) opened with EcoRI for experiments in mammalian cells. All obtained plasmids were sequenced verified by Sanger sequencing (complete plasmid sequences available upon request).

### Fibroblast cell culture

Fibroblasts from index patient 1 and her parents were obtained using a punch biopsy according to standard procedures, upon informed consent (IRB approval MEC-2017-341). Fibroblasts from the parents of index patient 2 and 3 were also obtained upon informed consent at McMaster Children’s Hospital. All fibroblasts were cultured in standard DMEM medium supplemented with 15% Fetal calf serum, MEM Non-Essential amino acids (Sigma), 100 U/ml penicillin and 100 µg/ml streptomycin, as done previously^108^, in routine humidified cell culture incubators at 20% O2. Fibroblast cell lines were transfected using Lipofectamine 3000 (Invitrogen) with the indicated plasmid constructs. All the cell lines used in this report were regularly checked for the presence of mycoplasma and were negative during all experiments.

### Genome engineering in human embryonic stem cells

H9 human embryonic stem cells were cultured as previously described^107,109^. In short, cells were maintained on feeder free conditions in mTeSR-1 medium (STEMCELL technologies) on Matrigel (Corning) coated culture dishes. To engineer the patient specific UGP2 mutation by homologous recombination^110^, ESC were transfected using Lipofectamine 3000 with a plasmid expressing eSpCas9-t2a-GFP (a kind gift of Feng Zhang) and a gRNA targeting the *UGP2* gene (see **Supplementary Table 8** for the sequence), together with a 60 bp single stranded oligonucleotide (ssODN) homology template encoding the patient mutation (synthesized at IDT). To increase the stability of the ssODN and therefore homologous recombination efficiency, the first two 5’ and 3’ nucleotides were synthesized using phosphorothiorate bonds^111^. 48 hours post transfection, GFP expressing cells were sorted, and 6000 single GFP-positive cells were plated on a Matrigel coated 6-well plate in the presence of 10µM ROCK-inhibitor (Y27632, Millipore). After approximately 10 days, single colonies where manually picked, expanded and genotyped using Sanger sequencing (see **Supplementary Table 8** for primer sequences). As a by-product of non-homologous end joining, knock-out clones were identified which showed a single nucleotide A insertion at position 42 of *UGP2* transcript 1 (chr2:64083462_64083463insA), leading to an out of frame transcript and a premature termination of the protein at amino acid position 47 (D15Rfs*33). Western blotting confirmed the absence of all UGP2 protein in knock-out clones and the loss of the short UGP2 isoform in clones with the patient mutation. To produce a stable rescue cell line, ESC cells were transfected as previously described with the pPyCAG-IRES-puro plasmid expressing either the long WT or mutant UGP2 isoform. After 48 hours, the population of cells with the transgene integration was selected with 1µg/ml puromycin. Engineered ESC clones had a normal colony morphology and pluripotency factor expression.

### Patient specific Induced pluripotent stem cell generation

Patient fibroblast cell lines were reprogrammed using the CytoTune™-iPS 2.0 Sendai Reprogramming Kit (Thermo Scientific, A16517) expressing the reprogramming factors OCT4, SOX2, KLF4 and C-MYC on matrigel coated cell culture plates, upon informed consent (IRB approval MEC-2017-341). After approximately 4-5 weeks, emerging colonies were manually picked and expanded. Multiple clones were assessed for their karyotype, pluripotency factor expression and three lineage differentiation potential (Stem Cell Technologies, #05230), following the routine procedures of the Erasmus MC iPS Cell facility, as previously described^108^. Sanger sequencing was used to verify the genotype of each obtained iPSC line. We used three validated clones for each individual in our experiments.

### Neural stem cell differentiation

Pluripotent cells were differentiated in neural stem cells (NSCs), using a modified dual SMAD inhibition protocol^112^. In short, 18000 cells/cm^2^ were plated on matrigel coated cell culture dishes in mTeSR-1 medium in the presence of 10µM Y27632. When cells reached 90% confluency, the medium was switched to differentiation medium (KnockOut DMEM (Gibco), 15% KnockOut serum replacement (Gibco), 2mM L-glutamine (Gibco), MEM Non-Essential amino acids (Sigma), 0.1 mM β-mercaptoethanol, 100U/ml penicillin and 100 µg/ml streptomycin) supplemented with 2µM A 83-01 (Tocris) and 2µM Dorsomorphin (Sigma-Aldrich). At day 6, medium was changed to an equal ratio of differentiation medium and NSC medium (KnockOut DMEM-F12 (Gibco), 2mM L-glutamine (Gibco), 20ng/ml bFGF (Peprotech), 20ng/ml EGF (Peprotech), 2% StemPro Neural supplement (Gibco), 100U/ml penicillin and 100µg/ml streptomycin) supplemented with 2µM A 83-01 (Tocris) and 2µM Dorsomorphin (Sigma-Aldrich). At day 10, cells were passaged (NSC p=0) using Accutase (Sigma) and maintained in NSC medium. We used commercially available H9-derived NSCs (Gibco) as a control (a kind gift of Raymond Poot, Rotterdam).

### Other stem cell differentiation experiments

ESCs were differentiated into hematopoietic stem cells and cardiomyocyte using commercially available STEMCELL technologies kits (STEMdiff Hematopoietic kit #05310, STEMdiff Cardiomyocyte differentiation kit #05010) according to manufacturer’s instructions. Cells were finally harvested and lysed with TRI reagent to isolate RNA for further RT-qPCR analysis.

### RNA-sequencing and data analysis

For RNA-seq on blood derived patient RNA, peripheral blood was obtained from index patient 1 and her parents, collected in PAX tubes and RNA was isolated following standard diagnostic procedures in the diagnostics unit of the Erasmus MC Clinical Genetics department. RNA-seq occurred in a diagnostic setting, and sequencing was performed at GenomeScan (Leiden, The Netherlands). For RNA-seq of *in vitro* cultured cell lines, RNA was obtained from 6-well cultures using TRI reagent, and further purified using column purification (Qiagen, #74204). mRNA capture, library prep including barcoding and sequencing on an Illumina HiSeq2500 machine were performed according to standard procedures of the Erasmus MC Biomics facility. Approximately 20 million reads were obtained per sample. For the cell line experiments, two independent H9 wild type cultures, two independent knock-out clones harboring the same homozygous *UGP2* genetic alteration and two independent clones harboring the patient homozygous *UGP2* mutation were used. Each cell line was sequenced in two technical replicates at ESC state and differentiated NSC state (at passage 5). FASTQ files obtained after de-multiplexing of single-end, 50 bp sequencing reads were trimmed by removing possible adapters using Cutadapt after quality control checks on raw data using the FastQC tool. Trimmed reads were aligned to the human genome (hg38) using the HISAT2 aligner^113^. To produce Genome Browser Tracks, aligned reads were converted to bedgraph using bedtools genomecov, after which the bedGraphToBigWig tool from the UCSC Genome Browser was used to create a bigwig file. Aligned reads were counted for each gene using htseq-count^114^ and GenomicFeatures^115^ was used to determine the gene length by merging all non-overlapping exons per gene from the Homo_sapiens.GRCh38.92.gtf file (Ensemble). Differential gene expression and RPKM (Reads Per Kilobase per Million) values were calculated using edgeR^116^ after removing low expressed genes and normalizing data. The threshold for significant differences in the gene expression was FDR < 0.05. To obtain a list of ESC and NSC reference genes used in Supplementary Figure 6F, we retrieved genes annotated in the following GO terms using GSEA/MSigDB web site v7.0: GO_FOREBRAIN_NEURON_DEVELOPMENT (GO:0021884), GO_CEREBRAL_CORTEX_DEVELOPMENT (GO:0021987), GO_NEURAL_TUBE_DEVELOPMENT (GO:0021915), BHATTACHARYA_EMBRYONIC_STEM_CELL (PMID: 15070671) and BENPORATH_NOS_TARGETS (PMID: 18443585).

### Functional enrichment analysis

Metascape^117^, g:profiler^118^ and Enrichr^119^ were used to assess functional enrichment of differential expressed genes. **Supplementary Table 4** reports all outputs in LogP, log(q-value) and Adjusted p-value (q-value) for Metascape and g:profiler, and in p-value, Adjusted p-value (q-value) and combined-score (which is the estimation of significance based on the combination of Fisher’s exact test p-value and z-score deviation from the expected rank) for Enrichr. All tools were used with default parameters and whole genome set as background.

### Genome-wide homology search

To make a genome-wide list of transcripts sharing a similar structure as UGP2 transcripts, 42976 transcripts from 21522 genes (Human genes GRCh38.p12) were extracted using BioMart of Ensembl (biomaRt R package). 11056 out of 21522 genes had only 1 transcript and the remaining 31920 transcripts from 10466 genes were selected, the protein sequences were obtained with biomaRt R package and homology analysis was performed using the NCBI’s blastp (formatting option: - outfmt=6) command line. We grouped longest and shorter transcript based on coding sequence length and only kept those that matched a pairwise homology comparison between the longest and the shorter transcript with the following criteria: complete 100 percent identity, without any gap and mismatch, and starting ATG codon of shortest transcript being part of the longest transcript(s). 1766 genes meet these criteria. We then filtered these genes for published essential genes^6^, leaving us with 1197 genes. Using BioMart (Attributes: Phenotype description and Study external reference) of Ensembl we then evaluated the probability that these genes were implicated in disease and identified 850 genes that did not have an association with disease phenotype/OMIM number. Of those, 247 genes encoded proteins of which the shorter isoform differed less than 50 amino acids from the longer isoform. We chose this arbitrary threshold to exclude those genes where both isoforms could encode proteins differing largely in size and might therefore encode functionally completely differing proteins (although we cannot exclude that this will also hold true for some of the genes in our selection).

### Differential isoform expression in fetal tissues

Publically available RNA-seq data from various fetal tissue samples (**Supplementary Table 2**) were analyzed using the same workflow as described for the RNA-seq data analysis above. To determine differential isoform expression in these tissues, we calculated a ratio between the unique exon(s) of the shortest and longest transcript for each gene and assessed its variability across different fetal tissue samples. The number of reads for each unique exon of a transcript was calculated by mapping aligned RNA-seq reads against the unique exon coordinate using bedtools multicov. The longest and shortest transcripts were separated and the transcript ratio (number of counts of shortest transcript / (number of counts of shortest transcript + number of counts of longest transcript)) for each gene was obtained from the average reads of RNA-seq samples per tissue. 382 genes out of 1197 genes showed high variability across different samples (defined as a difference between highest and lowest ratio > 0.5), 277 of those high variable genes were not associated with a disease phenotype/OMIM number and of these 83 genes had a length less than 50 amino acids (a subset of the 247 genes with no OMIM and length less than 50 amino acids)

### Haplotype Analysis

The 30 MB region surrounding UGP2 was extracted from exome sequencing VCF files to include both common and rare polymorphisms. Variants were filtered for a minimum depth of coverage of at least 10 reads and a genotype quality of at least 50. The filtered variants, were then used as input in PLINK (v1.07) with the following settings:

--homozyg-snp 5
--homozyg-kb 100
--homozyg-gap 10000
--homozyg-window-het 0

ROH around the *UGP2* variant were identified in all 5 probands examined. The minimum ROH in common between all samples was a 5Mb region at chr2: 60679942-65667235. We note that targeted sequencing leads to uneven SNP density, so the shared ROH may, in fact, be larger or smaller. Next, we used recombination maps from deCODE to estimate the size of the region in centiMorgans (cM). We then used the region size in cM to estimate the time to event in generations using methods previously described^120^.

### qPCR analysis

RNA was obtained using TRI reagent, and cDNA prepared using iSCRIPT cDNA Synthesis Kit according to manufacturer’s instructions. qPCR was performed using iTaq universal SYBR Green Supermix in a CFX96RTS thermal cycler (Bio-Rad). **Supplementary Table 8** summarizes all primers used in this study. Relative gene expression was determined following the ΔΔct method. To calculate the ratio of the short isoform, we performed absolute quantification as previously described^121^. Briefly, we performed qPCR on known copy numbers, ranging from 10^3 to 10^8 copies, of a plasmid containing the short UGP2 isoform (5’ UTR included) using primers detecting specifically either the total or the short isoform. After plotting the log copy number versus the ct, we obtained a standard curve that we used to extrapolate the copy number of the unknown samples. To test for significance, we used Student’s T-test and considered p<0.05 as significant.

### Western blotting

Proteins were extracted with NE buffer (20mM Hepes pH 7.6, 1.5mM MgCl2, 350mM KCl, 0.2mM EDTA and 20% glycerol) supplemented with 0.5% NP40, 0.5mM DTT, cOmplete Protease Inhibitor Cocktail (Roche) and 150U/ml benzonase Protein concentration was determined by BCA (Pierce) and 20-50µg of proteins were loaded onto a 4–15% Criterion TGX gel (Bio-Rad). Proteins were then transferred to a nitrocellulose membrane using the Trans-Blot Turbo Transfer System (Bio-Rad). The membrane was blocked in 5% milk in PBST and subsequently incubated overnight at 4°C with primary antibody diluted in milk. After PBST washes, the membrane was incubated 1 hour at RT with the secondary antibody and imaged with an Odyssey CLX scanning system (Li-Cor). Band intensities were quantified using Image Studio (Li-cor). Antibodies used were: Ms-α-UGP2 (sc-514174) 1:250; Ms-α-Vinculin (sc-59803) 1:10000; Gt-α-actin (sc-1616) 1:500; Ms-α-LAMP2 (H4B4) 1:200; IRDye 800CW Goat anti-Mouse (926-32210) 1:5000; IRDye 680 Donkey anti-Goat (926-32224) 1:5000.

### Zebrafish disease modelling

Animal experiments were approved by the Animal Experimentation Committee at Erasmus MC, Rotterdam. Zebrafish embryos and larvae were kept at 28°C on a 14–10-hour light–dark cycle in 1 M HEPES buffered (pH 7.2) E3 medium (34.8 g NaCl, 1.6 g KCl, 5.8 g CaCl_2_ · 2H_2_O, 9.78 g MgCl_2_ · 6 H_2_O). For live imaging, the medium was changed at 1 dpf to E3 + 0.003% 1-phenyl 2-thiourea (PTU) to prevent pigmentation. *Ugp2a* and *ugp2b* were targeted by Cas9/gRNA RNP-complex as we did before^76^. Briefly, fertilized oocytes from a tgBAC(slc1a2b:Citrine)re01tg reporter line^76^ maintained on an TL background strain were obtained, and injected with Cas9 protein and crRNA and tracrRNA synthesized by IDT (Alt-R CRISPR-Cas9 System), targeting the open reading frame of zebrafish *ugp2a* and *ugp2b*. DNA was extracted from fin clips and used for genotyping using primers flanking the gRNA location (**Supplementary Table 8**) followed by sequencing. Mutants with a high level of out of frame indels in both genes were identified using TIDE^122^ and intercrossed to obtain germ line transmission. Upon re-genotyping, mutant zebrafish with the following mutations as indicated in **Figure 6** were selected and further intercrossed. In this study, we describe two new mutant fish lines containing deletions in *ugp2a* (*ugp2a*^Δ/Δ^) and ugp2b (*ugp2b*^Δ/Δ^): *ugp2a*^*re08/re08*^ containing a 37 bp deletion in exon 2 and *ugp2b*^*re09/re09*^ containing a 5 bp deletion in exon 2. Intravital imaging, and analysis of eye movement, was performed as previously described^76^. Briefly, zebrafish larvae anesthetized in tricaine were mounted in low melting point agarose containing tricaine and imaged using a Leica SP5 intravital imaging setup with a 20×/1.0 NA water-dipping lens. To assess the locomotor activity of zebrafish larvae from 3 to 5 dpf, locomotor activity assays were performed using an infrared camera system (DanioVision™ Observation chamber, Noldus) and using EthoVision® XT software (Noldus) as described^76^. Briefly, control (*n* = 24) and *ugp2a*^Δ/Δ^; *ugp2b*^Δ/Δ^ (*n* = 24) zebrafish larvae, in 48 well plates, were subjected to gradually increasing (to bright light) and decreasing light conditions (darkness) as in Kuil et al^76^. Distance traveled (mm) per second was measured. For 4-AP (Sigma) stimulation animals were treated with 4-AP dissolved in DMSO 30 minutes before the onset of the experiments. For these experiments locomotor activity was measured over 35 minutes, with the first 5 minutes going from dark to light, followed by 30 minutes under constant light exposure.

### Periodic acid-schiff (PAS) staining

ESCs or differentiated NSCs (wild type, KO, KI or rescue) were incubated under hypoxia conditions (3% O2) for 48 hours. Cells were fixed with 5.2% formaldehyde in ethanol, incubated 10 min with 1% Periodic acid, 15 min at 37°C with Schiff’s reagent (Merck) and 5 min with Hematoxylin solution (Klinipath) prior to air drying and mounting. Every step of the protocol is followed by a 10 minutes wash with tap water. Imaging occurred on an Olympus BX40 microscope. Images were acquired at a 100x magnification, and ImageJ software was used for quantification. For ESCs, we used a minimum of 20 images per genotype for the quantification, containing on average 20 cells each, calculating the percentage of PAS positive area. For NSCs, we imaged between 80 to 100 cells per genotype, counting the number of glycogen granules in the cytoplasm. We report the average of two independent experiments at 48 hours low oxygen.

### UGP2 enzymatic activity

The measurement of UGP2 enzyme activity was performed according to a modified GALT enzyme activity assay as described previously^123^. Frozen cell pellets were defrosted and homogenized on ice. 10 µl of each cell homogenate (around 0.5 mg protein/ml as established by BSA protein concentration determination) was pre-incubated with 10 µl of dithiothreitol (DDT) for 5 min at 25°C. 80 µl of a mixture of glucose-1-phosphate (final concentration 1 mM), UTP (0.2 mM), magnesium chloride (1 mM), glycine (125 mM) and Tris-HCl (pH8) (40 mM) was added and incubated for another 15 min at 25°C. The reaction was stopped by adding 150 µl of 3.3% perchloric acid. After 10 min on ice the mixture was centrifuged (10,000 rpm for 5 min at 4°C), the supernatant isolated and neutralized with ice cold 8 µl potassium carbonate for 10 min on ice. After centrifugation the supernatant was isolated and 1:1 diluted with eluent B (see below) after which the mixture was added to a MilliPore Amicon centrifugal filter unit. After centrifugation the supernatant was stored at - 20°C until use. The separation was performed by injection of 10 µl of the defrosted supernatant onto a HPLC system with UV/VIS detector (wave length 262 nm) equipped with a reversed phase Supelcosil LC-18-S 150 mm x 4.6 mm, particle size 5 µm, analytical column and Supelguard LC18S guard column (Sigma-Aldrich). During the experiments the temperature of the column was maintained at 25°C. The mobile phase consisted of eluent A (100% methanol) and eluent B (50 mM ammonium phosphate buffer pH7.0 and 4 mM tetrabutylammonium bisulphate). A gradient of 99% eluent B (0-20 min), 75% eluent B (20-30 min) and 99% eluent B (30-45 min) at a flow rate of 0.5 m/min was used. The reaction product UDP-glucose was quantified using a calibration curve with known concentrations of UDP-glucose. UGP2 activity was expressed as the amount of UDP-glucose formed per mg protein per min. Experiments were performed in duplicate and for every cell line two independently grown cell pellets were used.

### Immunostaining / Immunohistochemistry

For immuonofluorescence staining, cells were seeded on coverslips coated with 100µg/ml poly-D-lysine (Sigma) overnight. For ESC, coverslips were further coated with Matrigel (Corning) for one hour at 37°C. When cells reached about 70% confluency, they were fixed with 4% PFA for 15 min at RT. Cells were then permeabilized with 0.5% triton in PBS, incubated one hour in blocking solution (3% BSA in PBS) and then overnight at 4°C with the primary antibody diluted in blocking solution. The following day the coverslips were incubated one hour at room temperature in the dark with a Cy3-conjugated secondary antibody and mounted using ProLong Gold antifade reagent with DAPI (Invitrogen) to counterstain the nuclei. Images were acquired with a ZEISS Axio Imager M2 using a 63X objective.

### Data availability

RNA-Seq of *in vitro* studies are publicly available through the National Center for Biotechnology Information (NCBI) Gene Expression Omnibus (GEO) under accession number GSE137129. A token for reviewer access is present in the supplement. Due to privacy regulations and consent, raw RNA-seq data from patient blood cannot be made available. To retrieve tissue wide expression levels of *UGP2*, the GTEx Portal was accessed on 16/07/2019 (https://gtexportal.org/home/). RNA-seq data from various tissues were downloaded from various publications^48-51^. All publically available data that were re-analyzed here are summarized in **Supplementary Table 2**.

## Figure Legends

**Supplementary Figure 1, related to Figure 1:**
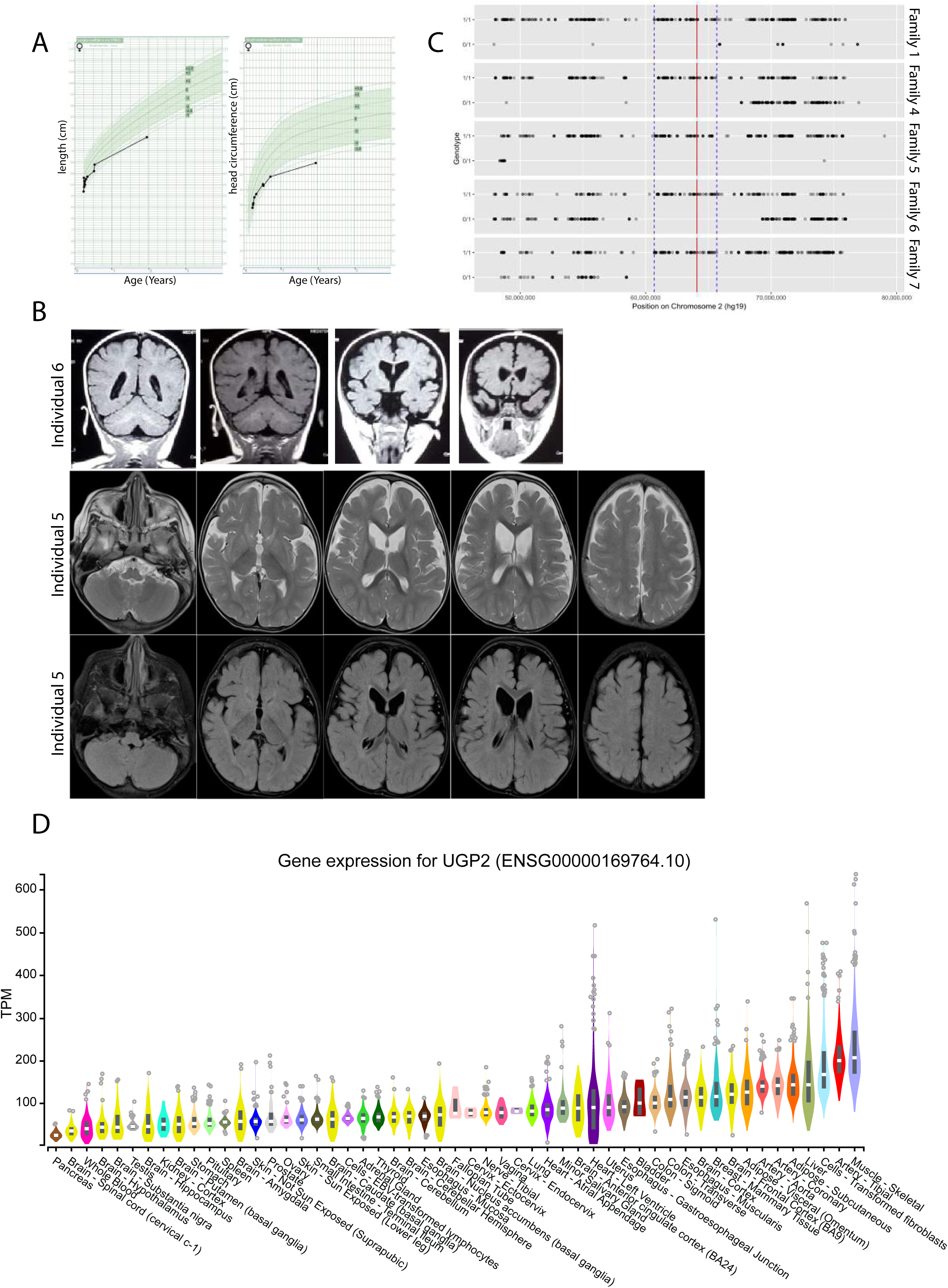
A) Growth chart from individual 1 for length (left) and head circumference (right) in cm. Reference chart from the Dutch population are used (TNO) and regions between −2 and + 2 SD are shaded. B) MRI studies of individual 5 (at the age of 12 month) and individual 6, showing global brain atrophy. C) ROH comparison between affected individuals from family 1, 4, 5, 6 and 7, carrying the homozygous chr2:64083454A>G mutation. The red line indicates the UGP2 variant, and the blue lines demark the shared ROH region between the individuals (chr2:60679942-65667235). D) Violin plots showing distribution of gene expression (in TPM) amongst samples from the GTEx portal^35^ for tissues and cell lines. Samples are sorted with the highest median TPM on the right. Outliers are indicated by dots.

**Supplementary Figure 2, related to Figure 2:**
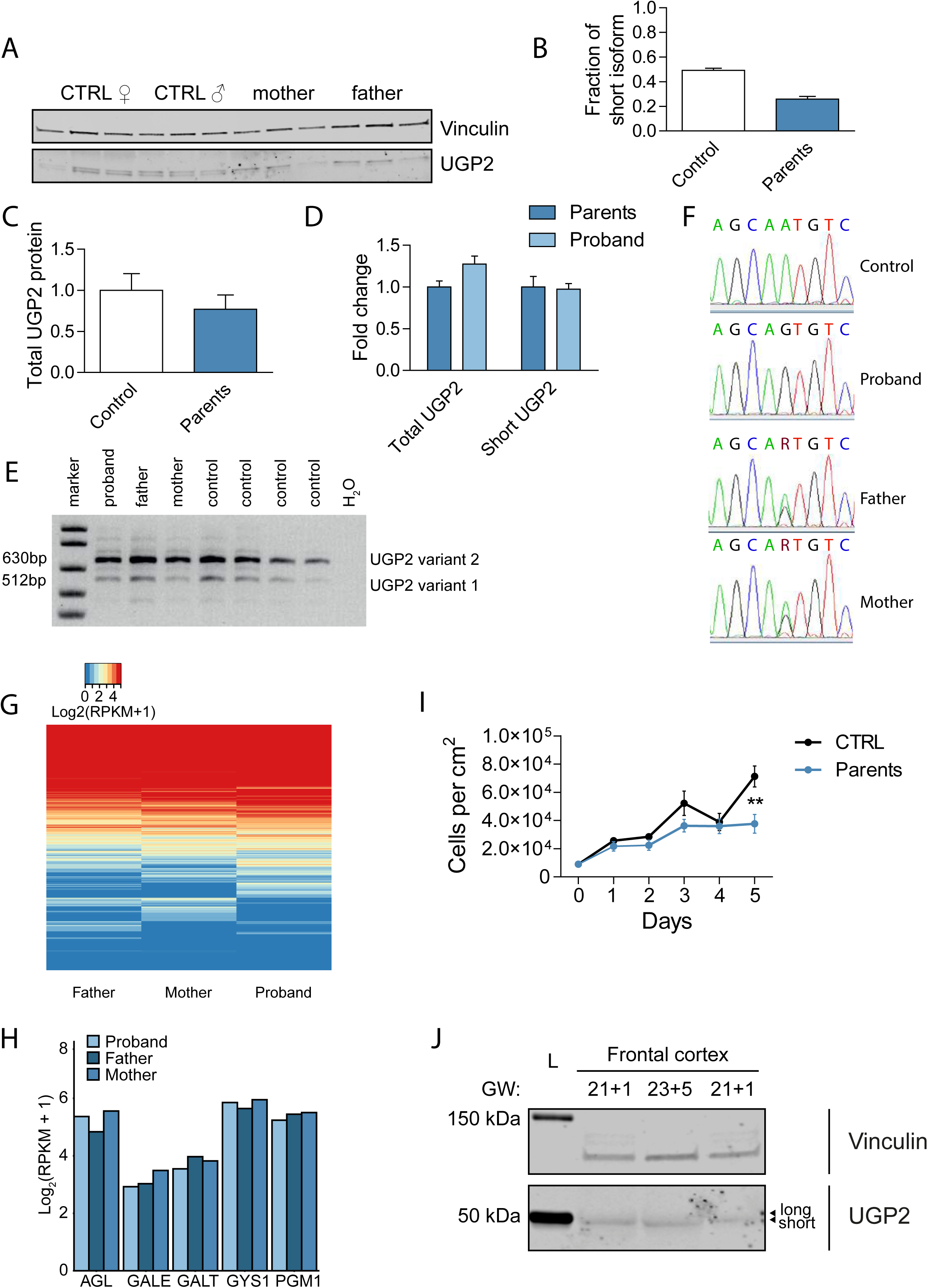
A) Western blotting of cellular extracts derived from control fibroblasts or fibroblasts obtained from heterozygous parents of family 2, detecting the house keeping control vinculin or UGP2. Note the two separated isoforms of UGP2 that have a similar intensity in wild type cells. The shorter isoform shows reduced expression in fibroblasts from heterozygous parents. B) Quantification of the fraction of the short UGP2 protein isoform compared to total UGP2 expression in control, and heterozygous fibroblasts from family 2, as determined in three independent experiments. Error bars represent SEM. C) Western blot quantification of total UGP2 protein levels, as determined by the relative expression to the housekeeping control vinculin. Bar graph showing the results from three independent experiments. Error bars represent SEM; no significant differences between control and parent samples, unpaired t-test, two-tailed. D) qRT-PCR analysis of total *UGP2* or the short isoform in fibroblast from heterozygous parents or homozygous proband from family 1, normalized for the housekeeping control *TBP*. The mean fold change compared to heterozygous parents of two biological replicates and two technical replicates is shown; error bars represent SEM no significant differences between control and parent samples, unpaired t-test, two-tailed. E) Multiplex RT-PCR detecting relative expression of *UGP2* isoform 1 and isoform 2 in peripheral blood from family 1 and unrelated wild type controls. F) Sanger sequencing of RT-PCR products from E), showing the expression of the homozygous and heterozygous chr2:64083454A>G *UGP2* variant in the index proband, her parents and an unrelated control. G) Heat map showing genome-wide gene expression levels (in log2(RPKM+1)) in peripheral blood from heterozygous parents and homozygous proband from family 1. H) Gene expression levels (in log2(RPKM+1)) from RNA-seq in peripheral blood for a selected number of genes involved in metabolism. I) Cell proliferation experiment of fibroblast from heterozygous parents from family 2 and wild type controls, during a 5 days period. Error bars represent SEM, **= p<0.01, unpaired t-test, two-tailed. J) Western blotting detecting UGP2 in human frontal cortex from week 21 and 23 of gestation, showing the virtual absence of the long isoform expression in fetal brain. Vinculin is used as a housekeeping control.

**Supplementary Figure 3, related to Figure 4:**
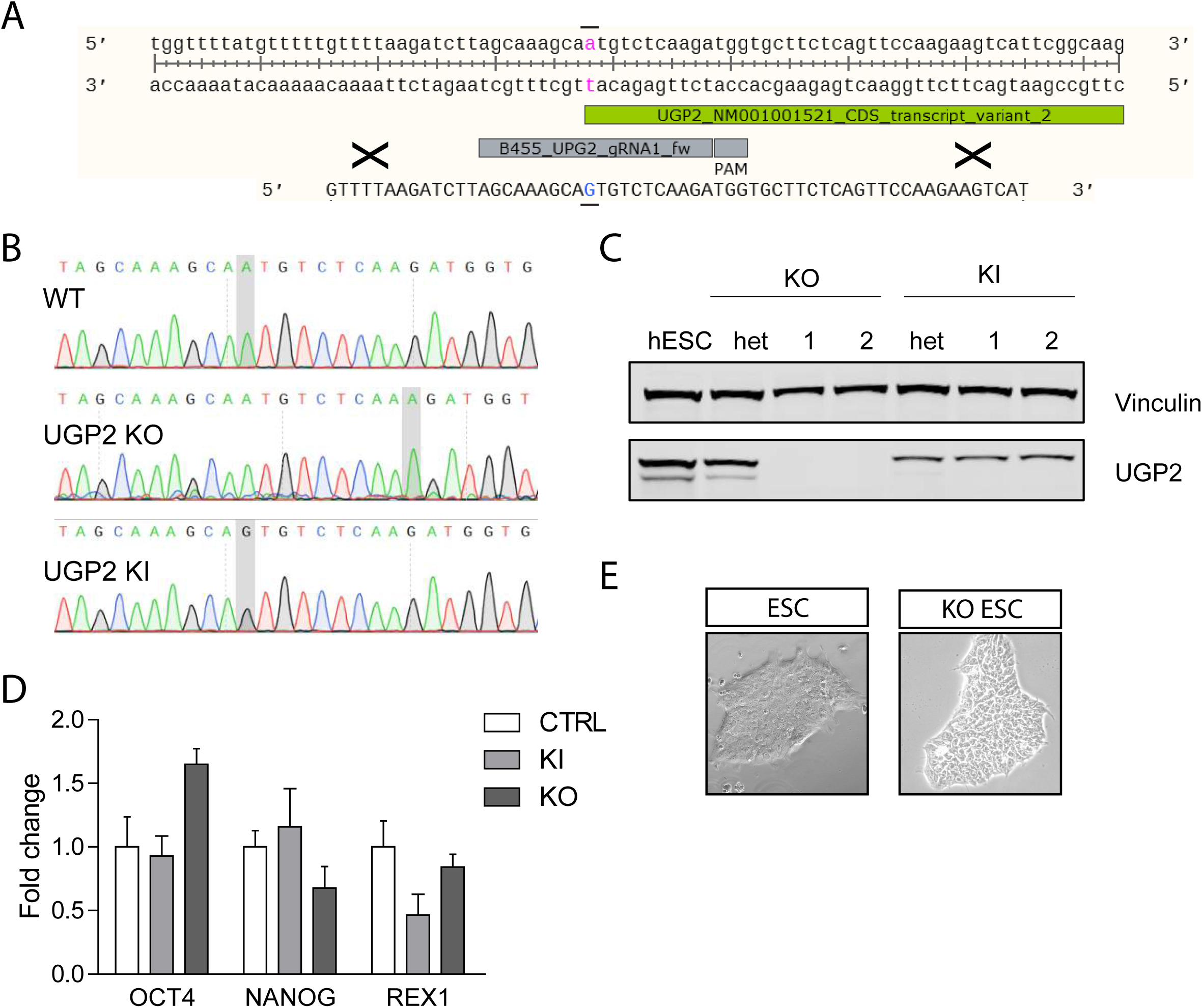
generation of mutant *UGP2* H9 cell lines A) Nucleotide sequence encompassing the ATG of *UGP2* transcript isoform 2. Indicated are the coding sequence, the location of the gRNA, PAM sequence and ssODN used to introduce the C.1A>G, p.? mutation. B) Sanger sequencing traces of part of the UGP2 gene from wild type, *UGP2* knock-out (KO) and *UGP2* knock-in H9 ESCs (KI). The A at the start of the coding sequence of UGP2 isoform 2 (short isoform) is highlighted. The homozygous insertion of an additional A in knockout and the mutation into a G in knock-in cells are indicated. C) Western blot detecting UGP2 and vinculin in wild type ESC, heterozygous and homozygous knockout and knock-in ESCs, as indicated. Note the complete loss of UGP2 in KO cells, and the loss of the short isoform in KI cells. D) RT-qPCR detecting the pluripotency factors OCT4, NANOG and REX1 in H9 wild type, UGP2 knock-in (KI) and *UGP2* knock-out (KO) ESCs, normalized for the house keeping control *TBP*. Mean fold change compared to wild type of two biological replicates and three technical replicates is shown; error bars represent SEM, *= p<0.05, unpaired t-test, two-tailed. E) Bright field image of a representative ESC colony from wild type parental and *UGP2* KO ESCs.

**Supplementary Figure 4, related to Figure 4.**
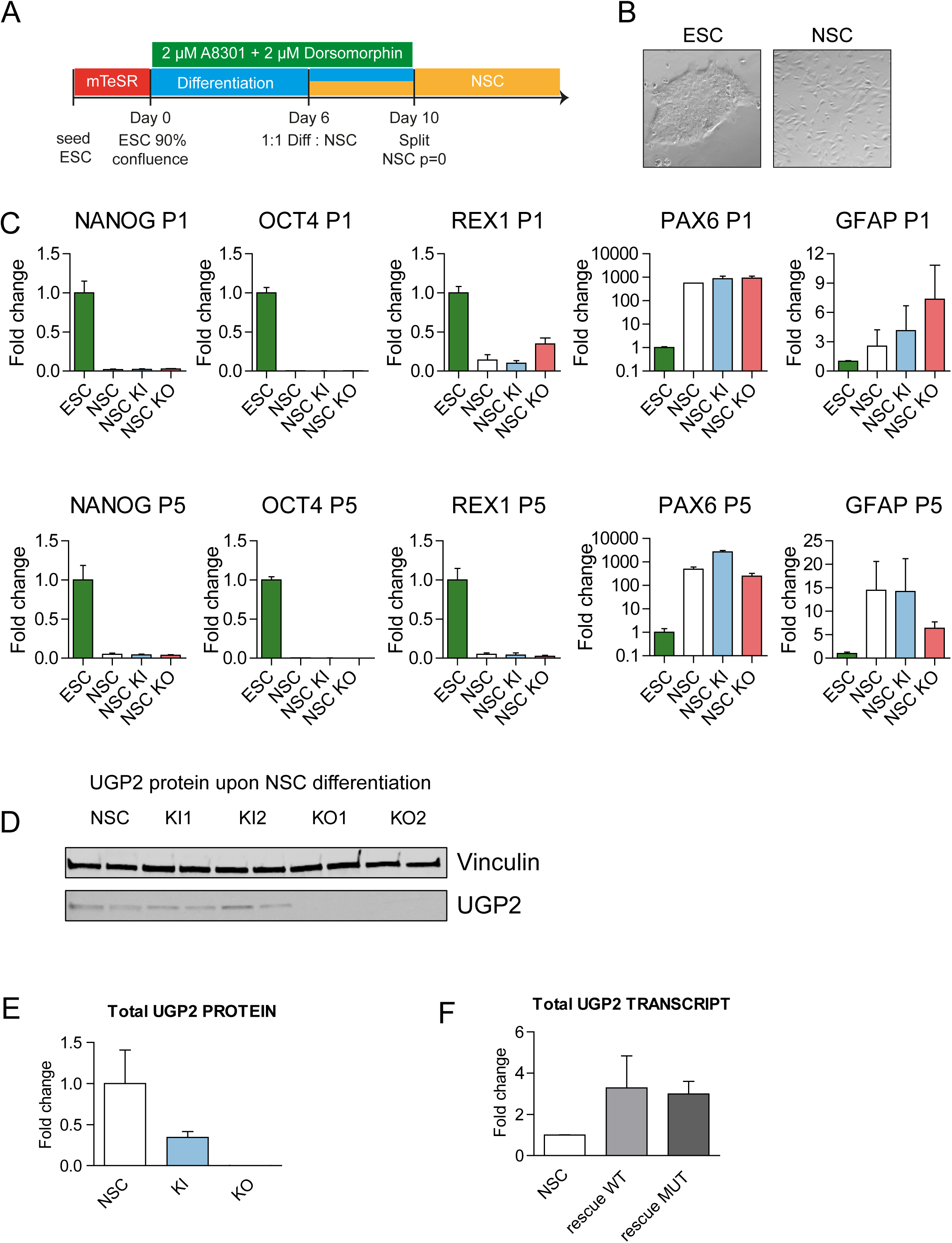
NSC differentiation. A) Schematic drawing of the differentiation procedure, see online methods for details. B) Bright field image showing representative pictures from ESCs and differentiated NSCs. C) qRT-PCR analysis for pluripotency markers (*NANOG, OCT4* (*POU5F1*), *REX1*) and genes expressed in NSCs (*PAX6, GFAP*) in WT, UGP2 KO and KI differentiated NSCs at p1 and p5. Mean fold change compared to wild type of two biological replicates and two technical replicates is shown; error bars represent SEM. D) Western blotting showing UGP2 expression in WT, UGP2 KI and KO differentiated NSCs. Vinculin is used as a housekeeping control. E) Quantification of total UGP2 protein levels by Western blot, as determined by the relative expression to the housekeeping control vinculin. Bar graph showing the results from two independent experiments; error bars represent SEM. F) qRT-PCR analysis of *UGP2* in NSCs or KO NSCs rescued with either the long wild type or long mutant UGP2 isoform. Mean fold change compared to wild type is shown for two biological replicates and three technical replicates; error bars represent SEM.

**Supplementary Figure 5:**
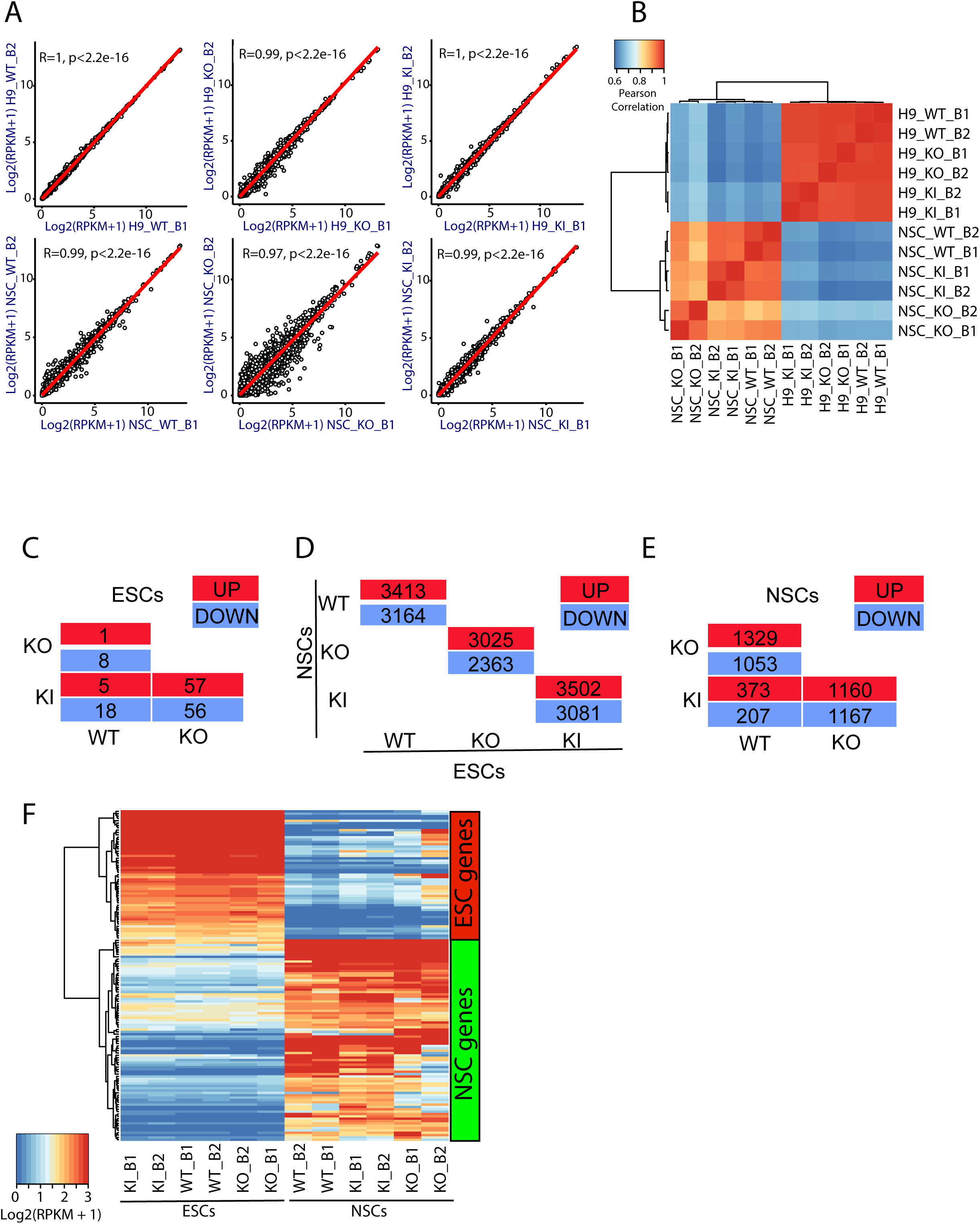
RNA-seq, related to Figure 4 A) Scatter plot showing the pair wise correlation between biological replicates. B) Heat map displaying Pearson correlation between biological replicates. C) Table summarizing up- (FDR<0.05 and LogFC>1) and down regulated (FDR<0.05 and LogFC<- 1) genes in WT, KO and KI ESCs. D) Table summarizing up- (FDR<0.05 and LogFC>1) and down regulated (FDR<0.05 and LogFC<-1) genes in WT, KO and KI ESC upon differentiation in NSCs. E) Table summarizing up- (FDR<0.05 and LogFC>1) and down regulated (FDR<0.05 and LogFC<-1) genes in WT, KO and KI NSCs. F) Heat map visualizing gene expression (in log2(RPKM+1)) and clustering of WT, KO and KI ESCs and NSCs, for a panel of ESC and NSC specific genes (see methods)

**Supplementary Figure 6, related to Figure 4:**
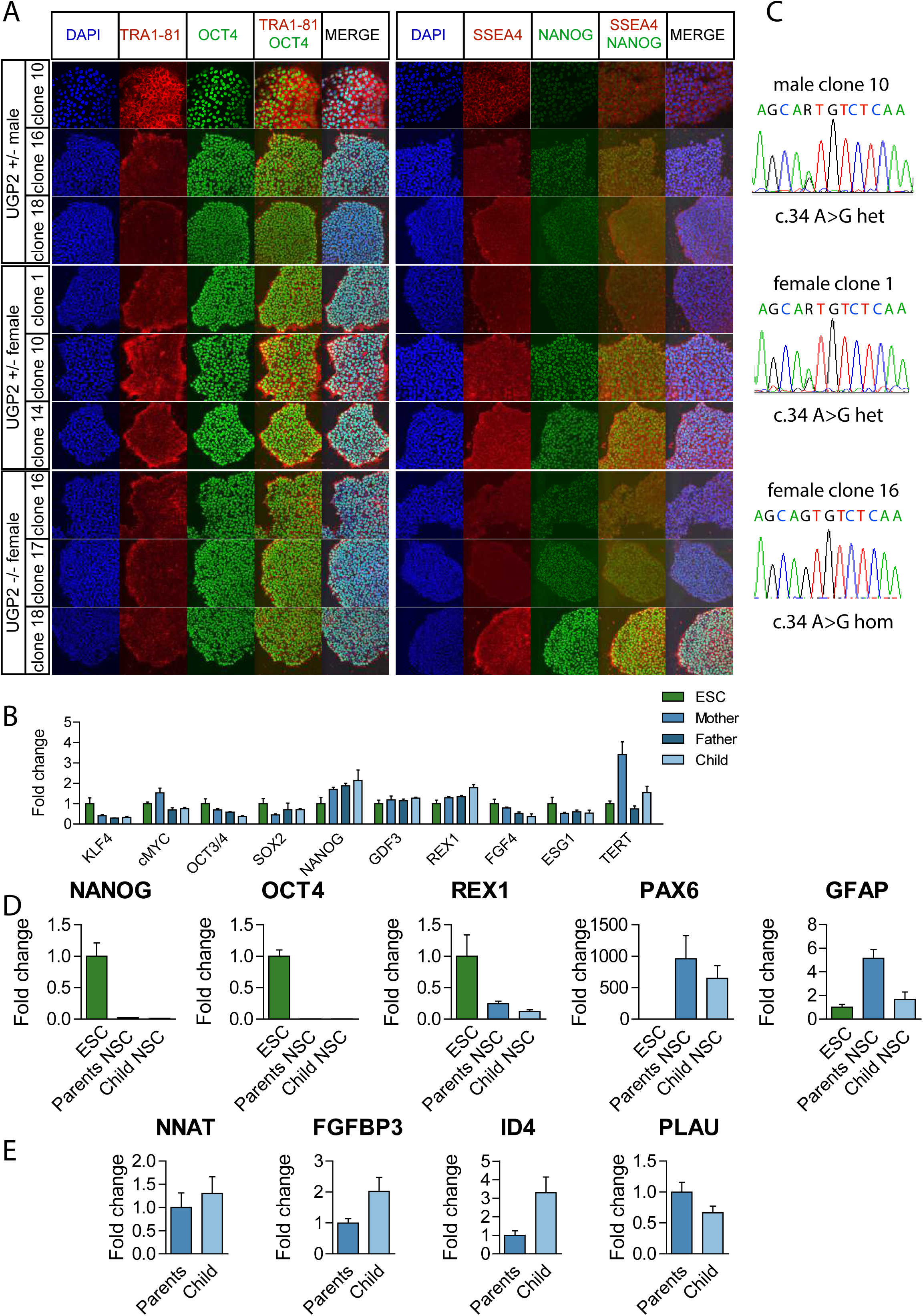
UGP2 mutant induced pluripotent stem cells A) Immunofluorescence of iPSC clones used in this study derived from Family 1 (three clones per individual) showing iPSC colonies stained for the pluripotency markers TRA1-81 (red) and OCT4 (green) (left panel) or SSEA4 (red) and NANOG (green) (right panel). Nuclei are stained with DAPI (blue). B) qRT-PCR expression analysis for the indicated pluripotency associated genes in 4 wild type control human embryonic stem cell lines and the iPSCs derived from family 1. Mean fold change compared to human embryonic stem cells of three biological replicates (e.g. individual clones from A) and three technical replicates is shown; error bars represent SEM. No statistically significant differences were found, unpaired t-test, two-tailed. C) Sanger sequencing of representative iPSC clones confirming the presence of the chr2:64083454A>G *UGP2* mutation in a heterozygous state in clones derived from parents and homozygous state in clones derived from the affected child. D) qRT-PCR PCR expression analysis upon differentiation for pluripotency (*NANOG, OCT4* (*POUF51*), *REX1*) and NSC markers (*PAX6, GFAP*), for H9 ESC control and heterozygous and homozygous iPSCs derived from family 1. Mean fold change compared to human embryonic stem cells of three biological replicates (e.g. individual clones from A)and two technical replicates is shown; normalized to *TBP*; error bars represent SEM. E) qRT-PCR expression analysis in iPSC-derived NSCs for genes that showed differential expression in RNA-seq experiments, e.g. *NNAT, FGFBP3, ID4* and *PLAU*. Mean fold change for cells obtained from the affected child compared to cells obtained from its parents (set to 1) of three biological replicates (e.g. individual clones from A) and two technical replicates is shown; normalized to *TBP*; error bars represent SEM.

**Supplementary Figure 7: related to figure 5:**
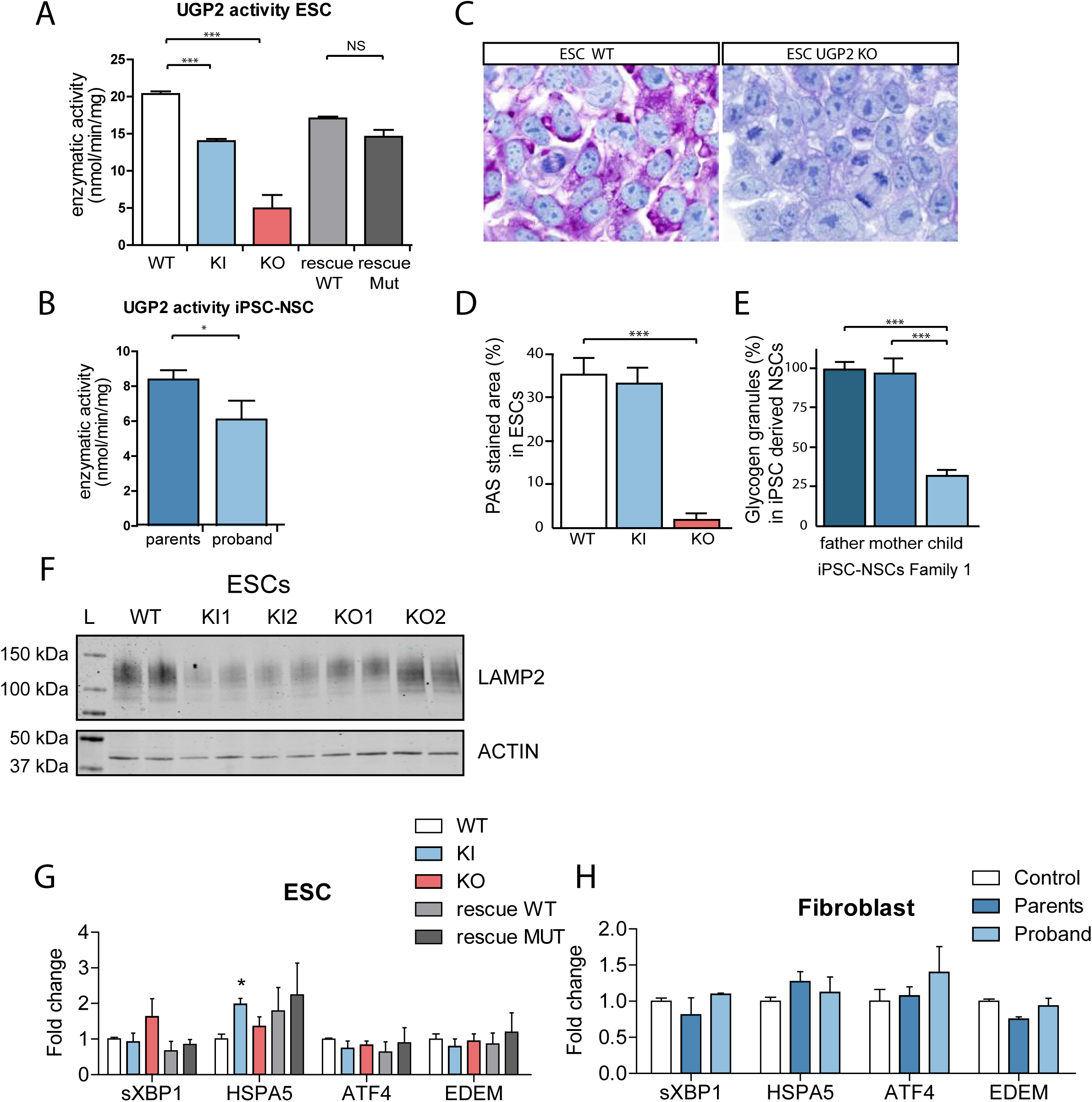
A) UGP2 enzymatic activity in WT, UGP2 KI, KO and KO ESCs rescued with wildtype isoform 1 or mutant Met12Val isoform 1 of UGP2. Plotted is the mean from two replicate experiments, error bar is SEM. ***=p<0.001, unpaired t-test, two-tailed. B) UGP2 enzymatic activity in iPSC derived NSCs from family 1. Plotted is the mean from two replicate experiments, measuring each the results for the three clones for each individual, error bar is SEM. *=p<0.05; unpaired t-test, two-tailed. C) PAS staining in WT and UGP2 KO ESCs. Nuclei are counterstained with hematoxylin (blue). D) Quantification of the PAS stained area in WT, KI and KO ESCs. Shown is the average PAS positive area per genotype from two biological replicates, each stained in two experiments; error bars are SD. ***=p<0.001, unpaired t-test, two-tailed. E) Glycogen granules detected by PAS staining in iPSC-derived NSCs from family 1 after 48 hours culture under low-oxygen conditions. Number of granules for paternal cell line are set at 100%. Average of three biological and two technical replicates per genotype, with each n=80-100 cells counted. Error bars represent SD, ***=p<0.001, unpaired t-test, two-tailed. F) Western blotting detecting LAMP2 (upper panel) and the house keeping control actin (lower panel) in cellular extracts from ESCs, that are WT, UGP2 KI, or KO. Compare to Figure 5D. G) qRT-PCR expression analysis for UPR marker genes (spliced *XBP1, HSPA5, ATF4* and *EDEM*) in WT, UGP2 KI, KO and rescue ESCs. Shown is the mean fold change for the indicated genes compared to wild type, normalized for the housekeeping gene *TBP*. Results of two biological and three technical replicates are plotted from two experiments. Error bars represent SEM; *= p<0.05, unpaired t-test, two-tailed). H) qRT-PCR expression analysis for UPR marker genes (spliced *XBP1, HSPA5, ATF4* and *EDEM*) in in primary fibroblasts from family 1. Shown is the mean fold change for the indicated genes compared to wild type, normalized for the housekeeping gene *TBP*. Results of two experiments with each three technical replicates are plotted. Error bars represent SEM; *= p<0.05, unpaired t-test, two-tailed.

**Supplementary Table 1**: Extended clinical characteristics of 11 patients with homozygous UGP2 variants

**Supplementary Table 2:** RNA-seq data used in this study

**Supplementary Table 3:** differentially expressed genes

**Supplementary Table 4:** enrichment analysis

**Supplementary Table 5:** UGP2 variants in *gnomAD*

**Supplementary Table 6:** genome-wide homology search results

**Supplementary Table 7:** *gnomAD* data of 247 disease candidate genes

**Supplementary Table 8**: Oligonucleotides used in this study

**Supplemental Movie 1**: wild type zebrafish eye movements

**Supplemental Movie 2**: Ugp2a/b double mutant zebrafish eye movements

### Supplementary Case Reports

**Supplementary Note**: additional example of an isoform start codon alteration of an essential gene

